# Population specific bottlenecks inflated differentiation measures of Louisiana black bear and negate subspecific status

**DOI:** 10.64898/2026.02.18.706691

**Authors:** Emily E. Puckett, Heather R. Clendenin

## Abstract

Significant debate has revolved around the delimitation of subspecific boundaries relative to conservation policy, and specifically how best to maximize limited resources. The conservation of subspecies captures intraspecific genetic diversity and aids in the long-term preservation of adaptive potential. Here, we evaluate patterns of neutral and adaptive genomic variation across the eastern lineage of the American black bear. We specifically assess the relative impact of phylogeographic history and local adaptation on differentiation of subpopulations in Louisiana, which were federally protected as a subspecies from 1992-2016. Despite high values of genetic differentiation (Fst >0.127) of these focal populations, we show that serial founder events during range expansion into eastern North America and multiple bottlenecks drove patterns of diversity within Louisiana. While we attribute initial population divergence between Louisiana subpopulations to east-west shifts of the Mississippi River between 6.2 – 2.7kya, drift accelerated following bottlenecks that were likely due to indigenous societies’ cultural and land-use changes and later to impacts of European fur traders. We further show that local adaptation has had a smaller impact (4.6%) than phylogeography (30.1%) on the distribution of genomic variation across this lineage. The strongest drivers of adaptive variation include mean annual temperature and monthly precipitation variation, where northern populations have substantial derived variation. Our genomic assessment, in conjunction with weak phenotypic data, does not support the continued recognition of *Ursus americanus luteolus* as a subspecies of American black bear. Continued genetic conservation efforts should focus on maintaining or increasing diversity, while supporting ongoing successes of demographic recovery.

## INTRODUCTION

While advances in applying phylogenetic data at the species level has improved clarity in taxonomic description, lower levels of taxonomic characterization have lagged behind (Gippoliti & Amori, 2007). The American black bear (*Ursus americanus*) is an example of this discrepancy; despite being a charismatic and well-studied species, their subspecific designations are an artifact of historical taxonomic practices that do not meet the criteria for contemporary delimitation (Kitchener et al., 2020; Puckett, Etter, Johnson, & Eggert, 2015). Currently, 16 subspecies are recognized (Lariviere, 2001), while contemporary genomic analyses suggest two to three lineages are present (Puckett et al., 2015).

Despite genuine scientific debate on the validity and best practices for subspecific delimitation, these trinomials are used to make practical conservation and management decisions (Haig et al., 2006).

Subspecies represent unique evolutionary trajectories within species and are distinguished by diagnosable phenotypes, embodying adaptive genetic diversity fine-tuned for ecological variation (Patten, 2015; B. L. Taylor et al., 2017). As such, targeting conservation efforts at infraspecific taxonomic units aims to prevent loss of evolutionary potential within species. Prioritizing subspecies for conservation also ensures that any potential losses of genetic diversity among populations may be buoyed by less imperiled and closely related populations elsewhere (Phillimore & Owens, 2006). However, since the U.S. Endangered Species Act (ESA) funding is finite, correct taxonomy is important for prioritizing available resources. Incorrect identification of subspecies can result in lack of protection for cryptic taxa when unique groups are not recognized, and conversely can result in unnecessary expenditures for taxa with over-split groups (Patten, 2015). Due to these consequences of inaccurate taxonomic assignment, it is important for taxonomists and managers to be in consensus regarding subspecific criteria and designations.

While American black bears are listed as “least concern” on the IUCN Red List (Garshelis, Scheick, Doan-Crider, Beecham, & Obbard, 2016), there are endangered populations within their Mexican range and small, isolated populations within the United States and Canada that may be threatened with extirpation. The latter group includes the four subpopulations within Louisiana (LA), designated as *U. a. luteolus* and known colloquially as the Louisiana black bear. *U. a. luteolus* is the only subspecies of American black bear to have been listed for protections under the ESA. These bears, especially along the Louisiana coast, are regarded as uniquely adapted to the heat and humidity characteristic of the region. Subpopulations have been of concern since the 1950s when roughly 80-120 bears remained in Louisiana. Efforts to bolster low population size, and increase genetic diversity, resulted in the translocation of 161 bears from Minnesota into the Lower Mississippi Alluvial Valley (LMAV) between 1964-1967 (Laufenberg et al., 2016). Specifically, 30 animals were moved into the Tensas River Basin, and the remaining 131 into the Upper Atchafalaya River Basin (UARB); the latter is considered a successful translocation (Laufenberg et al., 2016). Additional subpopulations were established via translocations within the LMAV in the 1980s (Laufenberg et al., 2016). Collectively, these demographic rescue efforts led to increases in the number of bears that met management benchmarks for census population criteria (Murphy et al., 2018). Evidence of population size recovery and genetic exchange between subpopulations (Laufenberg et al., 2016) was sufficient to merit the end of federal protections under the ESA, which lasted from 1992-2016 (USFWS, 1992, 2016). However, in such small, isolated populations, loss of individuals has a greater overall impact than in populations with a larger census size. As a result of habitat fragmentation and increased mortality across the interceding landscape matrix, gene flow is low or effectively nonexistent between some subpopulations (Murphy et al., 2018).

In this study, we use genomic data to respond to questions raised in previous microsatellite-based analyses with respect to the evolutionary uniqueness of the Louisiana black bear and investigate historical demography, timing of differentiation, and the distribution of adaptive, neutral, and deleterious variation across populations of bears in eastern North America. While we find differentiation statistics in Louisiana bears equivalent to species-level differences in other taxa, these patterns do not appear to be caused by natural selection for local adaptation to the environment. Instead, genetic differentiation has been driven by neutral forces such as range expansion, landscape barrier formation, habitat alteration, and multiple anthropogenically-driven bottlenecks. Additionally, we investigate the footprint of the translocation efforts mid-20^th^ century and evaluate the genetic consequences of bears introduced to LMAV from the Great Lakes region. Overall, we find that, while demographic recovery was achieved, genetic recovery will require continued management efforts.

## METHODS

### Sample Collection

We partnered with state and federal wildlife agencies, and university partners in North America which collected tissue (i.e., ear punch, muscle, or blood) from bears that were hunter harvested, vehicle killed, or live animals captured for other research studies or management purposes. Samples were collected from 2011–2023 and metadata included locality and field called sex (Puckett et al., 2023; Puckett et al., 2015).

### Whole Genome Sequencing and Variant Calling

We sequenced 173 *U. americanus* animals from the eastern lineage (Table S1) to an average depth of 3.7x. Specifically, we followed the manufacturer’s instructions for the Illumina DNA Prep Kit except that we diluted the reactions to 20% volume and started with 20ng of genomic DNA. Pooled libraries of 24 samples were sequenced on one lane of an Illumina 4000 at Novogene (Davis, CA). An additional 10 samples were sequenced to an average depth of 27.8x, where library preparation was not diluted. We mapped reads to the *U. americanus* reference genome (Srivastava et al., 2019) using BWA-MEM v0.7.17 with default parameters.

For population structure, ROH, and selection analyses, we imputed the low coverage data using a reference panel of American black bears from across the range (Puckett et al., 2023). Sites were called with GATK v.4 (McKenna et al., 2010), then imputed with GLIMPSE v.1 (Rubinacci, Ribeiro, Hofmeister, & Delaneau, 2021) as described previously (Puckett et al., 2023) but with the expanded sampling. As imputation only occurs for sites within the reference panel, this dataset had 7,369,660 biallelic SNPs prior to analysis specific filtering. Following imputation, the high coverage samples were merged into the data then phased with BEAGLE v5.4 (Browning, Zhou, & Browning, 2018).

As the imputation strategy limits the dataset to alleles within the reference panel, we decided on an alternative genotype likelihood (GL) framework to capture additional eastern lineage specific variation for the Stairway and deleterious allele analyses. GLs for individual populations or the entire dataset were calculated using ANGSD v0.941 (Korneliussen, Albrechtsen, & Nielsen, 2014). We first determined the ancestral alleles across the genome using the -doFasta 2 function within ANGSD. Specifically, we download one sample from each of: sloth (*U. ursinus*; SlB03), sun (*U. maylanus*; SuB13), Asian black (*U. thibetanus*; SRA DRR250459), and brown (*U. arctos*; SRA SRR 20985023) bears (Puckett et al., 2023; Zou et al., 2022). Samples were mapped to the American black bear reference genome, then .bam files were used as input into ANGSD (-doCounts 1 -snp_pval 0.01 -domaf 1 -domajorminor 1 -gl 2 -rmSNPs 1 -explode 1 -doFasta 2). Sites were only called if all four samples were observed (-minind 4) and the minimum depth of each individual at a site was at least 15 (-setMinDepthInd 15). The output fasta file with ancestral variants was used with the -anc flag in downstream GL estimations where we set the ancestral allele as the major allele (-doMajorMinor 5).

For Stairway plots (Liu & Fu, 2015), we output an unfolded one-dimensional site frequency spectra (SFS) per population. The American black bear reference genome and our estimates of ancestral alleles (see above) were provided as inputs, the SAMTOOLS genotype likelihood calculator (-GL 1), a minimum mapping and minimum quality score of 20 (-minMapQ 20 and - minQ 20), -C 50, and -baq 1 were included as parameters. For analyses of the entire eastern lineage, we utilized the reference and ancestral genomes, the above four parameters, then added: minimum depth per individual (-setMinDepthInd) of 1, maximum depth across samples (-maxDepth) of 2340, and minimum number of individuals observed per site (-minInd) of 92 (i.e., 50%). This resulted in 9,055,411 SNPs identified across scaffolds 2–37, and output as a VCF with GLs.

### Population Structure

Before assessing population structure, we filtered variants within the imputed dataset, using VCFTOOLS v0.1.16 (Danecek et al., 2011), to remove factors that may confound analyses. First, we masked all annotated genes along with 50kb of sequence upstream of the start codon and downstream from the stop codon. Second, we removed sites with a minor allele frequency <0.05, and for sites with >10% missing data. Next to removed linked sites, we thinned the dataset to one variant within non-overlapping 10kb windows. The final dataset for population structure contained 77,193 SNPs.

We evaluated population structure using both PCA and ancestry plots. The PCA was conducted within PLINK v1.9 (C. C. Chang et al., 2015). Ancestry was assessed using ADMIXTURE (Alexander, Novembre, & Lange, 2009), where 20 iterations of each cluster (K) value between 1–25 were run. The cross-validation error was calculated for each cluster and plotted to infer the best fitting model to the data.

To understand the pattern of population divergence, we ran TREEMIX v1.12 (Pickrell & Pritchard, 2012). To create an outgroup for this analysis, we first mapped three *U. arctos* animals from the Greater Yellowstone Ecosystem population (SRR20985035, SRR20985034, SRR20985023) to the *U. americanus* genome. We used ANGSD to call variants at the same sites as used in the population structure analysis, then merged the outgroup genotypes into the working VCF file. We used blocks of 1,000 SNPs and tested 0-9 migration edges. We assessed the proportion of variance and the residuals of the population tree and focused our interpretation on a population with one migration edge.

### Change in Population Size Through Time

Previous estimates of genetic diversity (Murphy et al., 2018) or changes in effective population size (N_E_) over time (Puckett, 2025) suggested that population declines may have occurred relative to post-colonial and/or LGM processes. We refine the patterns and timing of bottlenecks in four focal populations: Great Lakes [Minnesota and Michigan], Southeast [Appalachian Mountains], Coastal [Louisiana (LA)], and Tensas [LA]. Specifically, we input the SFS of each population into Stairway Plots (Liu & Fu, 2015). For each population, a stairway blueprint file was made with the number of chromosomes (2 * sample number), the SFS, and four break points (number of input chromosomes minus two, then multiplied by 0.25, 0.5, 0.75, or 1).

As Stairway plots rely on the coalescent for inference, they miss changes in N_E_ in generations nearest to the present; therefore, we additionally ran GONe (Santiago et al., 2020) to capture N_E_ within the last 100 generations. GONe uses patterns within the distribution of linkage disequilibrium (LD) to estimate recent N_E_. For five focal populations (Great Lakes, Southeast, Coastal, Tensas, and UARB where individuals with less than 90% ancestry assignment to UARB were removed so as not to bias for contemporary admixture), we exported a phased VCF for each population, then input into GONe using default parameters including a recombination rate of 0.05. We ran 50 iterations of the program which each drew 10k sites per scaffold for the estimation of LD. We calculated the median, and 2.5% and 97.5% confidence intervals from the repetitions in R before plotting. Initially run with the imputed dataset, we were concerned that population specific rare variants may bias estimates; therefore, we also exported the GLs for each population, then imputed and phased the GL data using BEAGLE. We did not observe differences in patterns between the two datasets and therefore report the one from our imputed dataset.

### Population Divergence Timing

To infer the pattern and timing of population divergence across the eastern lineage, we utilized MSMC-IM (Wang, Mathieson, O’Connell, & Schiffels, 2020). As input, we first estimated the change in effective population size (N_E_) through time on each of the high coverage samples using MSMC2 (Schiffels & Durbin, 2014). Data preparation, including the construction of genome-wide masks with SNPable (Li, 2009) was previously reported (2025); however, comparisons were made among four focal populations (Great Lakes, Southeast, Coastal, and Tensas). For all populations except Tensas, two samples (four haplotypes) were used, whereas only a single high-depth sample was available for Tensas, so only two haplotypes were available. Following MSMC2, the cross-coalescent results were input into MSMC-IM to estimate the patterns and timing of population divergence and the rate of gene flow over time. Output was converted to years and N_E_ using a mutation rate of 1×10^−8^ (Kumar & Subramanian, 2002) and a generation time of 6.5 years (Onorato, Hellgren, Van Den Bussche, & Doan-Crider, 2004).

### Inbreeding and Genetic Diversity

The extent of inbreeding across populations were estimated by identifying runs of homozygosity (ROH) in individual samples. We identified ROH using the sliding-window-based method in PLINK. We selected parameters to keep the scanning window small to detect short ROHs and avoid underestimating F_ROH_. These parameters included: homozyg-window-threshold 0.05, homozyg-window-missing 5, homozyg-window-het 5, homozyg-density 50, homozyg-het 1. We set the maximal gap to 500kb (homozyg-gap 500). Each scanning window had a minimum size of 100kb (homozyg-kb). The minimum number of SNPs per window (homozyg-window-snp) and the final ROH segment (homozyg-snp) were based on a species-specific estimate (Clendenin, Pollard, & Puckett, 2025) of the number of SNPs per ROH intended to eliminate artificially small ROH due to chance (Gorssen, Meyermans, Buys, & Janssens, 2020; Purfield, Berry, McParland, & Bradley, 2012). We did not prune by linkage disequilibrium nor minor allele frequency as this can result in missing long ROHs (Meyermans, Gorssen, Buys, & Janssens, 2020). We used the identified ROH to estimate F_ROH_, a genomic coefficient of inbreeding, for each individual. To compare relative timing of inbreeding across populations, we compared the number of ROHs per individual to the sum total length of ROHs in their genome (as longer ROHs indicate more recent consanguineous mating).

We used individual heterozygosity as a proxy for overall genetic diversity across populations. Observed heterozygosity (H_O_) was estimated genome-wide for each individual by finding the proportion of heterozygous sites compared to the total number of sites mapped in the genome using VCFTOOLS.

### Estimation of Deleterious Load

In a limited sample of high coverage genomes, our previous work showed that the high diversity Great Lakes region harbored more deleterious load than the low diversity subpopulations in Louisiana (Clendenin et al., 2025), consistent with population genetic theory relative to N_E_ and the impacts of bottlenecks (Clendenin et al., 2025). Yet, inbreeding depression has been suggested for the Louisiana populations (Murphy et al., 2018). Thus, we wanted to understand the impact of shared and private deleterious sites across eastern lineage American black bears.

Deleterious variation within and across populations was assessed by annotating and analyzing the GL and imputed VCFs. Using our 9M site dataset, we annotated each variant according to the model in SnpEff (Cingolani et al., 2012). SnpEff predicts deleterious variants within genic regions based on sequence annotation and variant categories. SnpEff annotations for variants include functional categories (sense, missense, stop-gained, etc.) and the expected impact upon gene function. SnpEff was used to identify variants in three categories: loss of function (LOF; including frameshifts, start-lost, and stop-gained), missense, and synonymous variants. To make the VCFs compatible with the SnpEff database, positional information was lifted over from the DNAzoo assembly to the NCBI assembly using HalLiftover (Hickey, Paten, Earl, Zerbino, & Haussler, 2013).

Relative load per population was assessed by computing the relative frequency of variants (R_XY_) by category (Do et al., 2015). Frequencies of variants in each category and each population were generated, and the Great Lakes, Tensas, and Coastal populations were compared against the Southeast group. Mean R_XY_ values, jackknife standard errors, and 95% confidence intervals were calculated across chromosome-excluded replicates.

To compare allele frequency distributions, 2dSFS were generated for each pair of the four focal populations (Great Lakes, Southeast, Coastal, and Tensas). ANGSD SFS estimates were generated using protocols similar to those described above, but with a sites file for each category containing the genomic location of each variant identified by SnpEff as LOF, missense, or sense.

Claims of recent admixture due to translocation of *U. a. americanus* into the UARB subpopulation were investigated through additional comparisons between the Great Lakes, the UARB, and the rest of the LMAV populations. We first removed putative F_1_ individuals from the UARB and remaining LMAV populations based on previously described ancestry assignments, then generated new ANGSD SFS estimates for each group. Using custom UNIX and R scripts, unique and shared variants across each grouping were identified and counted, variant frequencies were identified, and 2dSFS were generated for each pairwise comparison.

We tested for functional over-representation of both biological processes and molecular function using the PANTHER v19.0 web portal (Mi et al., 2005; Thomas et al., 2022). As our analysis was limited to scaffolds 2–37, we input a background gene list annotated from those scaffolds (n = 15,834) so as not to bias estimates.

### Geographic Patterns of Adaptation

Although eastern North America has less variable habitat types and altitudes when compared to western North America, we remained interested in which environmental factors may have driven local adaptation within the eastern lineage. We proportioned the amount of genomic variation explained by the environment and spatial structure of the landscape using redundancy analysis (RDA) (Forester, Lasky, Wagner, & Urban, 2018; Lasky et al., 2012), a type of genotype-environment association.

As input genomic data, we first removed samples from the UARB of Louisiana and the Ozark Mountains of Missouri, as both populations have histories of translocations from the Great Lakes region (Smith & Clark, 1994; D. F. Taylor, 1971). The recent admixture and/or population replacement (Murphy et al., 2018; Puckett et al., 2014) of the genomic signatures would act as a confounding factor within our analysis of selection to environmental gradients. We then removed loci with a MAF < 0.05, before thinning so that loci were not within 20kb of each other. The final dataset had 105,364 SNPs.

The second input into an RDA are environmental layers expected to impact the axes of selection. We downloaded BioClim variables 1-19 (Hijmans, Cameron, Parra, Jones, & Jarvis, 2005), WorldClim variables 20-35 (Sosa-Guillén, González, Pérez, Expósito, & Díaz, 2024), the humidity layer for August from BioClim+ (Brun, Zimmermann, Hari, Pellissier, & Karger, 2022), and an altitude layer, at a spatial extent of 30-arc seconds. All environmental layers were input into a GIS using ArcPro v2.3 (ESRI; Redlands, CA), then the cell value for each layer was exported at the coordinates for our samples across eastern North America.

To capture the spatial configuration of the landscape itself, we created a set of distance-based Moran’s eigenvector maps (dbMEMs). These are eigenvectors created from Moran’s I, a spatial statistic that identifies patterns of autocorrelation and is able to account for variation due to cardinal direction orientation (e.g., anisotropy). We followed the procedure outlined in Chang et al (2022); specifically, the mean longitude and latitude for each subpopulation (n = 13) was used to create a Gabriel graph network. Spatial weights were created for the network, then the inverse distance was calculated which served as input to create the dbMEM vectors using the *adespatial* R package (Dray, Legendre, & Peres-Neto, 2006). From our dataset, 12 orthogonal vectors were output. We used the forward selection method (forward.sel) to identify dbMEMs significantly (*P* < 0.05) associated with genomic variation, and all were retained.

We calculated the correlation between all environmental layers (n = 42) and dbMEM vectors (n = 13) using the correlation (cor) function in R (R Core Team, 2020), then removed one layer for any pair with an r^2^ > 0.70. We retained nine continuous variables: mean annual temperature (Bio01), temperature of the wettest quarter (Bio08), coefficient of variation of precipitation (Bio15), precipitation of the warmest quarter (Bio18), mean temperature of the driest month (Bio30), moisture seasonality (Bio31), potential evapotranspiration (Bio34), altitude, and August humidity. We tested three RDAs, one each for the spatial scale (with 12 significant dbMEM layers) and environmental variables (e.g., nine layers) alone, then a partial RDA where we accounted for spatial scale within an analysis of the environmental variables. For the partial RDA, we removed dbMEM3 due to correlation with environmental variables; thus, there were 11 dbMEMs and nine environmental variables in the analysis. ANOVAs were used to assess the significance of each variable within the full model using 1000 permutations. All nine environmental variables contributed to both the full and partial RDA and the contribution is reported as a precent.

### Selection and Diversification Scans

To generate the selective regions, we utilized three population genetic metrics meant to capture different aspects of differentiation and selection. Between populations, we used sliding *F_ST_*windows (i.e., six tests) to identify regions of the genome that had elevated differentiation. Estimates were calculated with VCFTOOLS with a 50kb window size and 50kb slide such that windows were non-overlapping. As single population tests, we calculated integrated haplotype score (iHS) (Voight, Kudaravalli, Wen, & Pritchard, 2006) and integrated haplotype homozygosity (iHH12) (Garud, Messer, Buzbas, & Petrov, 2015). iHS and iHH12 vary in their ability to detect hard and soft sweeps, respectively. Estimates were made from SELSCAN v2 (Szpiech, 2022; Szpiech & Hernandez, 2014). Given the bottleneck intensity within the two focal Louisiana populations and expectations for strong drift, we used conservative thresholds to identify genomic regions with possible selection. For all three tests the top 0.1% of sites or windows were retained from each population or comparison. Further, from the partial RDA, we identified SNPs within 3 standard deviations from the mean axis loading score for the first three axes, and associated each site with the environmental variable driving the loading. These sites were considered candidates under selection.

We intersected the positions or windows identified as being selected upon across the eastern lineage with the genome feature file of the reference genome using BEDTOOLS v2.31 (Quinlan & Hall, 2010). Specifically, we accepted annotations within a 10kb window. To test if selected genes were enriched for specific biological processes, we ran a GO term analysis in PANTHER as described above.

### Visualization

To aid in interpretation of our results, we digitized Figure 1 of Gouw and Autin (2008) within ArcPro which represents the fluvial geomorphology of the lower Mississippi River, then added dates to the river staging based on Saucier (1994).

**Figure 1.**
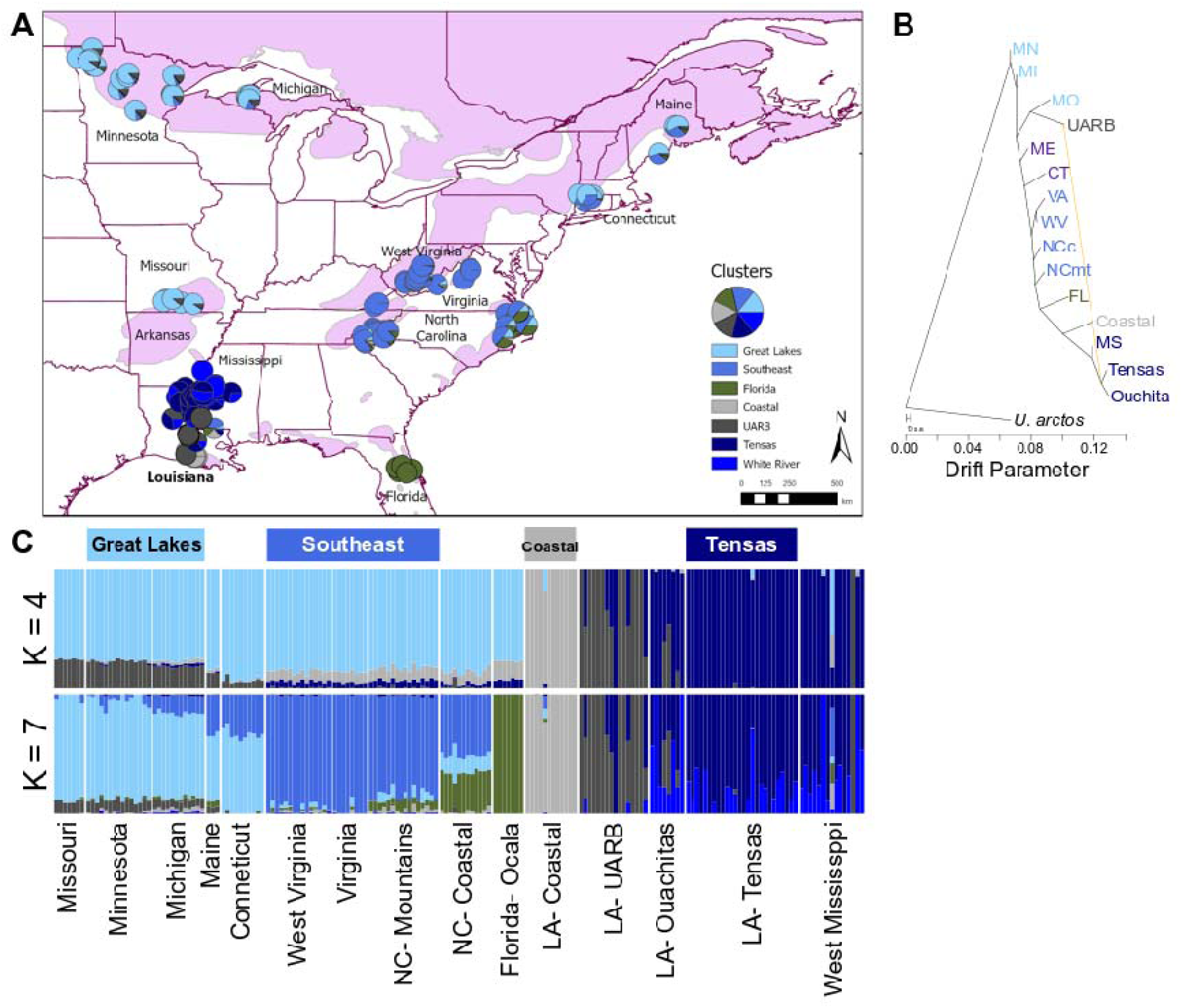
Geography of the eastern lineage of *Ursus americanus* (American black bear) highlighting the contemporary range of the species (violet) with USA state names of sampled regions denoted. (A) Map of the study area where pie charts represent ancestry proportions of seven clusters (light blue- Great Lakes; medium blue- Southeast; olive green- Florida; light grey- Coastal, Louisiana (LA); dark grey- Upper Atchafalaya River Basin (UARB), LA; navy- Tensas, LA; and royal blue- White River Basin, Arkansas). (B) A population tree of sampled areas created using TREEMIX with the inclusion of a 13% migration edge, where brown bear (*U. arctos*) rooted the tree. (C) Ancestry plots at both four and seven clusters (K) from ADMIXTURE. Samples included from four focal areas highlighted throughout the paper are denoted at the top.

To display the partial RDA model, we first created 5,000 random points (ArcPro, Create Random Points) within the *U. americanus* species distribution map, where no point was closer than 0.5 decimal degrees. We deleted points westward of the eastern edge of Manitoba, Canada, leaving 1,492 points to analyze. Next, we extracted the values from the eight BioClim and altitude layer using the Extract Multi Values to Points tool in ArcPro, then exported the coordinates and variables into a text file. This was imported into R where the coefficients for each variable from the nine RDA axes were applied to the data points. The predicted spatial values were smoothed with the Kriging function in ArcPro using a spherical semivariogram, and colors were applied at across nine Jenks natural breaks in each distribution for visualization.

## RESULTS

We sequenced 173 and 10 *U. americanus* animals to an average depth of 3.7x and 27.8x, respectively (Table S1). We mapped reads to the *U. americanus* reference genome (Srivastava et al., 2019) using BWA-MEM v0.7.17, called sites with GATK v.4 (McKenna et al., 2010), then imputed with GLIMPSE v.1 (Rubinacci et al., 2021). The dataset contained 7,369,660 biallelic SNPs prior to filtering.

All populations were previously placed within the eastern lineage of the American black bear (Puckett et al., 2015), and our first aim was to understand the spatial genetic structure of this lineage. The population tree from TREEMIX (Pickrell & Pritchard, 2012) (Figure 1B) shows a range expansion pattern beginning with the Great Lakes populations, expanding eastward towards Maine, dropping southwards, before a western expansion likely along the gulf coast, then finally northwards into the LMAV. The presence of bears in eastern Mexico as early as 16kya (Pedersen et al., 2021) suggests continued westward expansion past LMAV and approximate timing of bears in the southern range extent. Our best supported number of clusters was seven; however, we focus on four populations (Great Lakes, Southeast, Coastal, and Tensas; Figures 1, S1-S3) that contrast in latitude and demographic history, and then highlight unique signatures within Louisiana. While ancestry proportions vary among populations in the eastern lineage, we interpret this data as serial founder events along the phylogeographic path of range expansion, rather than as secondary contact.

Genetic differentiation between (sub)populations in the eastern lineage, excluding those in Louisiana, ranged from low to moderate *F_ST_*(0.019 – 0.139; Table S2). Differentiation elevated when populations within Louisiana were compared to the remainder of the lineage with *F_ST_* varying 0.127 – 0.295. Within Louisiana, differentiation between the Coastal and Tensas subpopulations was 0.282. These elevated values were consistent with increased drift observed in the population tree backbone.

### Eastern Lineage Range Expansion Initiated During the LGM

Given the phylogeographic pattern, we next aimed to estimate when population divergence initiated relative to the Last Glacial Maximum (LGM), which pushed eastern lineage populations into a Southeastern glacial refugia (Puckett et al., 2015). MSMC-IM (Wang et al., 2020) estimated initial divergence between the eastern and western lineages ∼138kya, where 50% of ancestry coalesced by 99.2kya (grey line in Figure 2A) (Puckett, 2025). From there, our results show that northern populations diverged earliest, followed by the Southeast (Figures 2A, S4). Within the Southeast, the Tensas and Coastal subpopulations of Louisiana began differentiating late in the LGM, with divergence accelerating around 5.5kya (Figure 2A) then reaching 50% coalescence ∼5.0kya. The timing of differentiation between these populations coincides with stage 3 of the Mississippi River (Saucier, 1994). At that time, the path of the river would have been significantly shifted to the west compared to the current flow (Figure 2C), thereby forming a barrier to movement across the landscape that would have been compounded by unpredictable flooding and landscape instability (Gouw & Autin, 2008). We further note a deep time spike in bidirectional gene flow between the Louisiana subpopulations ∼800kya (Figure S4) which may indicate a geographically isolated ancient admixture/hybridization event, or a coalescent artifact due to bottlenecks (see below).

**Figure 2.**
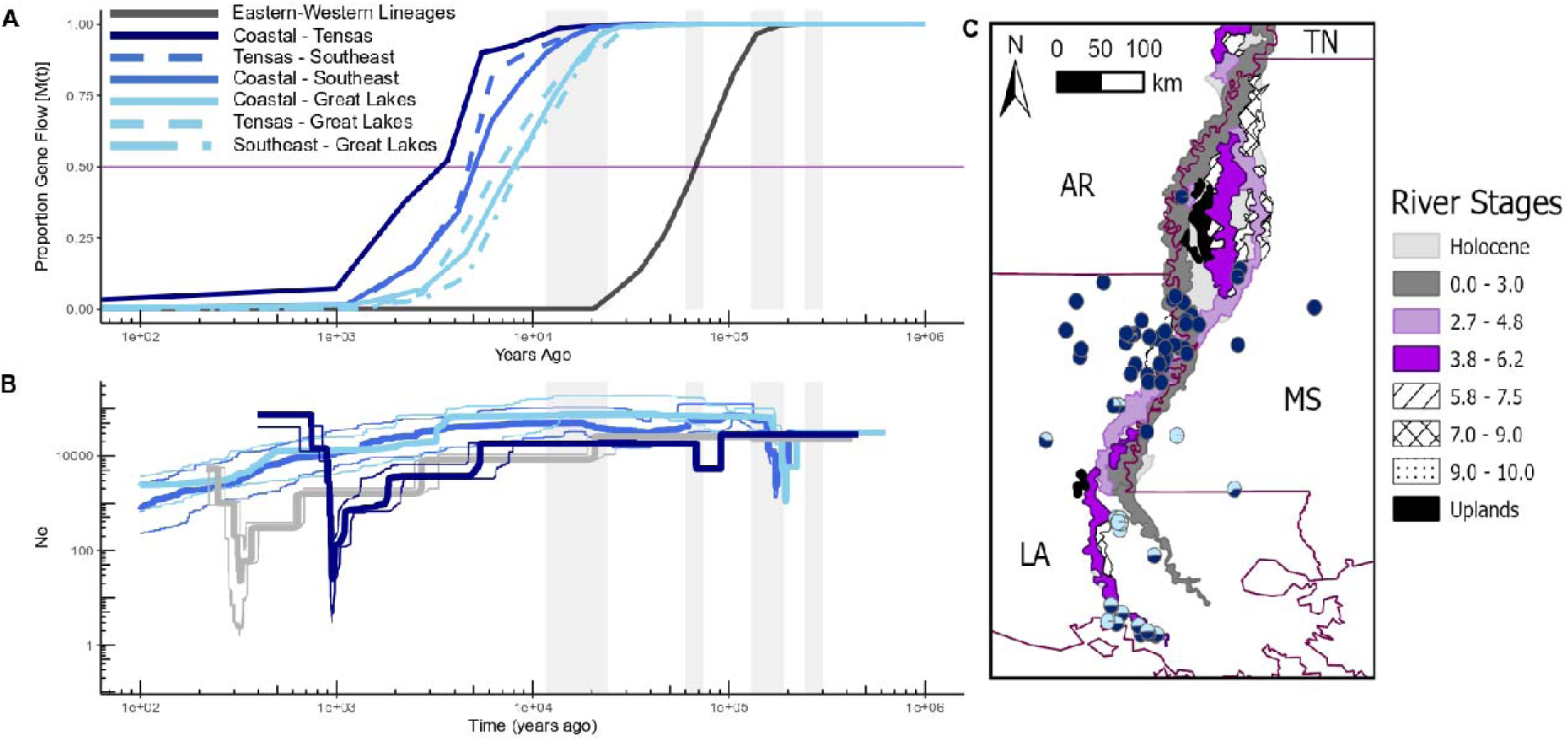
Demography of four populations within the eastern lineage of *Ursus americanus* (American black bears). (A) Change in the probability of divergence between two populations (Great Lakes and Coastal (light blue and solid; 50% coalescence 11.7kya); Great Lakes and Tensas (light blue and regular dash; 9.7kya); Great Lakes and Southeast (light blue and long-short dash; 12.5kya); Southeast and Coastal (medium blue and solid; 7.5kya); Southeast and Tensas (medium blue and regular dash; 6.9kya); Coastal and Tensas (navy and solid; 5.0kya); and the eastern and western lineages (dark grey; 99.2kya)). Estimates were generated from MSMC-IM and the probability of panmixia is displayed on the y-axis over time, where divergence estimates are taken at the 50% (violet line). Light grey background indicates glacial periods (left to right: Marine Isotope Stages 2, 4, 6, and 8). (B) Effective population size (N_E_) through time as estimated using SFS-based Stairway plots highlight unique and independent bottlenecks in the Tensas (navy) and Coastal (grey) populations of Louisiana. While declining N_E_ since the Last Glacial Maximum (i.e., MIS 2) were observed for the whole lineage, bottlenecks for the Southeast (medium blue) and Great Lakes (light blue) populations were only apparent in MIS 6 around the timing of the eastern lineage founder event. Thick lines indicate median estimates with thin lines highlighting the 2.5 – 97.5% confidence interval. (C) Detailed map of the Lower Mississippi Alluvial Valley (LMAV) with shifts in the river stage (i.e., barriers to dispersal) colored based upon the timing of their presence on the landscape in thousands of years ago (kya) (Gouw & Autin, 2008). The light and medium grey represent the current flood plain of the Mississippi River. Stages 2 (light violet) and 3 (bright violet) indicate barriers on the landscape over time due to shifts in fluvial geomorphology during relevant temporal periods of interest. Older stages are denoted by varying hashes. The contemporary USA state boundaries are denoted in maroon (LA- Louisiana; MS- Mississippi; AR- Arkansas; TN- Tennessee).

### Effective Population Size Declined Due to Both Climate and Pre- and Post-Colonial Anthropogenic Landscape Changes

While the ESA listing for Louisiana black bear cited habitat fragmentation and loss (USFWS, 1992) which often decrease effective population size (N_E_), we are also aware of additional species-level N_E_ declines since the LGM (Clendenin et al., 2025). Thus, we quantified change of N_E_ across both historical and recent timescales. Using Stairway Plots (Liu & Fu, 2015), we observed that N_E_ declined in all four focal populations after the LGM (Figure 2B). Beyond long-term declines, the Tensas and Coastal subpopulations of Louisiana each show extreme and population-specific bottlenecks. In Tensas, the bottleneck occurred 1kya (∼1020 AD) and reduced N_E_ to ∼3% of its pre-bottleneck size. The later bottleneck in the Coastal subpopulation occurred 0.33kya (∼1690 AD) and reduced population size to ∼6%.

Land-use changes in the 19^th^ and 20^th^ centuries are also believed to have driven population bottlenecks. We used the linkage-based GONe (Santiago et al., 2020) to estimate changes in N_E_ over the past 100 generations, during which these impacts would have taken effect. In Tensas, N_E_ declined sharply 30 generations ago (∼200 years), dropping from approximately 5,000 to 50 (Figure S5A). Declines in Coastal began between 25 – 22 generations ago, when N_E_ changed from 1,000 to a low of 50, then rebounded to 100. The Great Lakes and Southeast populations also underwent sharp declines 8 and 15 generations ago, respectively (Figure S5B). The Great Lakes N_E_ was estimated at 8,000 and declined to approximately 500 currently, whereas the Southeast started near 3,000 and has a contemporary N_E_ of 320, highlighting widespread declines in bear populations due to habitat loss and persecution.

### Louisiana Populations have High Inbreeding and Low Genetic Diversity but have Purged Deleterious Alleles

Southern populations of American black bears are characterized by small and isolated habitat patches, which is expected to increase inbreeding and reduce genetic diversity. To quantify the consequences of this isolation, we estimated individual genomic coefficients of inbreeding (F_ROH_) and genome-wide observed heterozygosity (H_O_). The proportion of the genome within a run of homozygosity (ROH) tract was lowest within the Great Lakes (mean F_ROH_ = 0.054, SD = 0.017; Figure 3A) and moderate within the Southeast (mean = 0.121, SD = 0.035). However, the southernmost and highly isolated populations had higher F_ROH_, including in Florida (mean = 0.262, SD = 0.043) and across the LMAV (mean = 0.361, SD = 0.100). The pattern in H_O_ was similar but mirrored, with the Great Lakes having the highest H_O_ (mean = 6.7 × 10^−4^, SD = 2.5 × 10^−5^; Figure 3B, Table S1), followed by the Southeast (mean = 5.8 × 10^−4^, SD = 3.1 × 10^−5^). Diversity declined in Florida (mean = 4.9 × 10^−4^, SD = 2.8 × 10^−5^), and was lowest among Louisiana bears (mean = 4.6 × 10^−4^, SD = 7.6 × 10^−5^).

**Figure 3.**
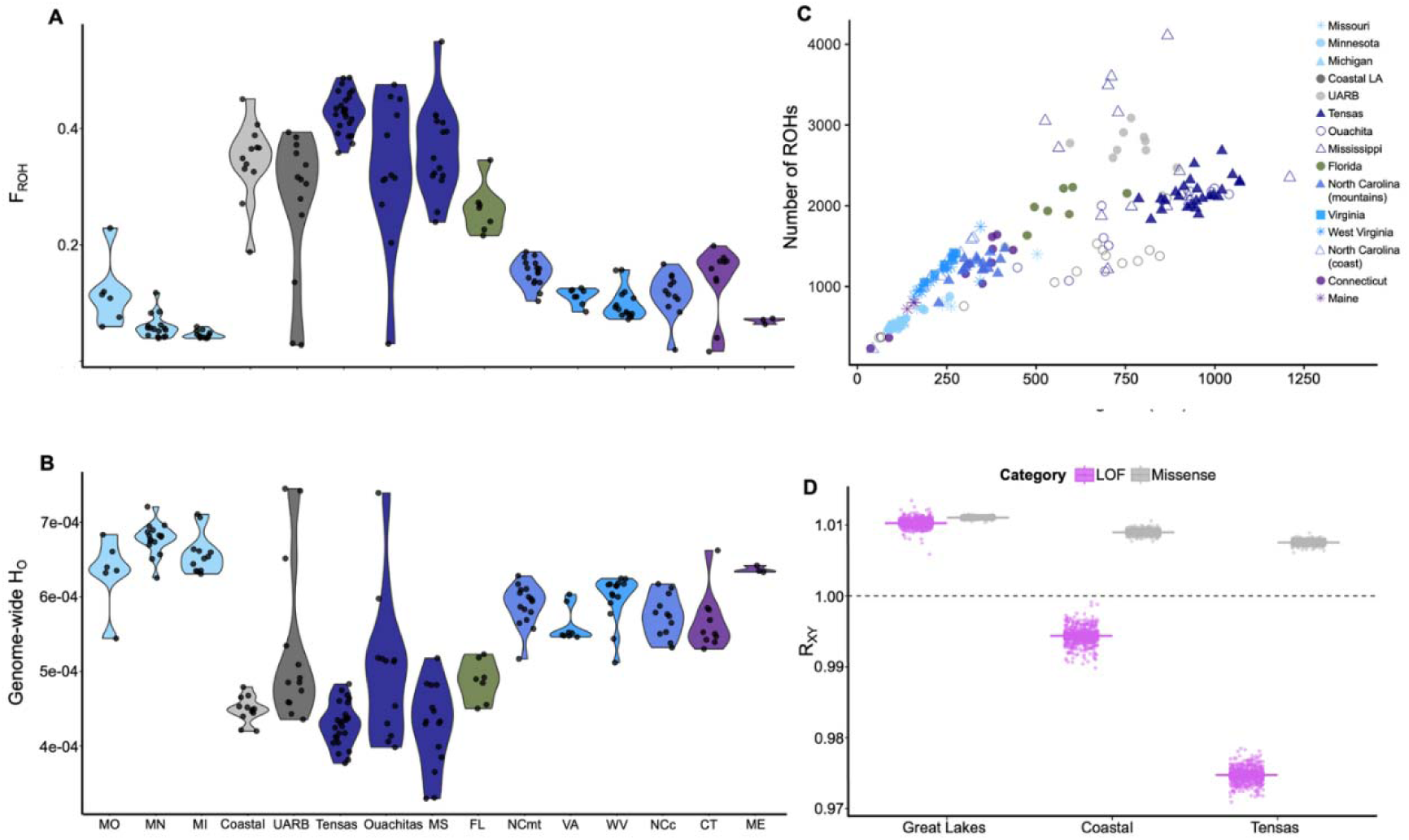
Diversity and deleterious load across the eastern lineage of *Ursus americanus* (American black bear). (A) The genomic coefficient of inbreeding, F_ROH_, by population. (B) Genome-wide observed heterozygosity (H_O_). (C) Number of runs of homozygosity (ROH) by the sum total length of ROHs across individual genomes. Right skewed points highlight individuals with recent inbreeding within their pedigree. (D) Relative frequency (R_XY_) for loss of function (LOF; violet) and missense (grey) variants calculated across the Great Lakes, Coastal, and Tensas populations compared to the Southeast as a baseline. Dashed line indicates R_XY_ = 1; variant categories with a frequency R_XY_ >1 indicate that the frequency of those mutations are greater frequency in the featured population than in the Southeast; R_XY_ <1 indicate the frequency of mutations in that category are lower in the featured population than in the Southeast. Solid bars represent the mean R_XY_ value, while individual points represent values from chromosome-excluded replicates used to generate confidence intervals.

We compared the frequency and timing of consanguineous matings between populations by looking at differences across individuals in the number and total length of ROH tracts across the genome (Figure 3C). ROH tracts are most abundant among small, inbred populations and at their longest immediately after consanguineous mating events, becoming shorter over subsequent generations as they are broken up by recombination. The lowest distribution and shortest total length of ROHs were found among individuals in the Great Lakes, along the northeastern coastline, and across the southeast (excluding Florida and Louisiana). Individuals in Florida and Coastal Louisiana fell along the same slope but with higher values in the Coastal group, suggesting that the timing of consanguineous matings may be similar but that the Coastal bears inbreed at a higher frequency. Louisiana individuals generally had greater total lengths of ROHs in the Tensas subpopulation and slightly fewer tracts there than in Coastal bears, suggesting relatively more overall inbreeding events and recent or ongoing consanguineous mating in Tensas.

The consequences of inbreeding vary by genetic load, which we assessed via annotation with SnpEff (Cingolani et al., 2012). We identified putatively deleterious variation across the eastern lineage, then compared the distribution of load among populations. We identified 24,420 sense, 18,489 missense, 996 nonsense, and nine start loss variants among the 183 samples (Table S3). Among those variants, 20 sense and 28 missense were fixed within the lineage. Genes with fixed derived variants may indicate early sites of selection to the unique climate of eastern North America; however, there were no over-represented GO terms among the genes for the fixed variants. We next generated pairwise 2dSFS plots among protein-coding variant types to understand patterns of allele sharing (Figure S6). Broadly, the Great Lakes and Southeast show signatures of population divergence, with segregating variants found largely at low frequencies within each population. In Louisiana, the Coastal population exhibited low to moderate allele sharing across the frequency spectrum with both Southeast and Great Lakes. This contrasted to Tensas, which shared most alleles with the Coastal population but otherwise had erratic allele frequency sharing patterns with the other three populations. We interpret this as the effects of random drift across the allele frequency spectrum.

Given our results indicating multiple factors that can impact genetic load (i.e., bottlenecks, decreased genome-wide diversity, and shifts in allele sharing), we next tested whether deleterious alleles had been purged from either Louisiana population. We calculated R_XY_ for both loss of function (LOF) and missense variants. We observed greater purging of LOF variants in Tensas compared to Coastal (R_XY_ values of 0.975 ± 3.0 × 10^−6^ and 0.994 ± 3.4 × 10^−6^, respectively; Figure 3D). Purging of missense variants was not detected (Tensas: 1.007 ± 1.2 × 10^−6^ and Coastal: 1.009 ± 1.3 × 10^−6^), which is consistent with population genetic theory regarding the accumulation of slightly deleterious alleles in populations with lower N_E_. Our estimates also demonstrate that the population that experienced an earlier bottleneck had sufficient time for purging highly deleterious load.

### Adaptation Variation is Limited within the Eastern Lineage

During the time when bears expanded their range and colonized eastern North America, the landscape would have been covered in diverse broad and/or needle leaf forested habitats. In contrast to habitats colonized by the western lineage of American black bears, eastern North America is considerably wetter and more humid. While we hypothesize lineage-wide adaptation to those two specific climatic variables, additional local adaptation is expected across the range relative to ecological variation.

We ran two full and one partial redundance analyses (RDA) to explain the proportion of genomic variation partitioned among the spatial configuration of the landscape and environmental variables, then identified specific environmental variables relevant for local adaptation. The first RDA used 12 distance-based Moran’s Eigenvectors Maps (dbMEMs; Figure S8) to account for the spatial configuration of the sampled populations, and had an adjusted r^2^ of 28.1%. This suggests that large amounts of spatial genomic variation are due to the phylogeographic pattern.

The second RDA included eight environmental variables and altitude, which explained 22.1% of the genomic variation, where all environmental variables were significant (Table S4). From this analysis, the first two RDA axes recapitulated the geographic pattern of the samples and showed distinct separation of Tensas from other populations (Figure S9A).

To decompose the effects of spatial configuration on environmental variation, we ran a partial RDA in which 11 dbMEMs were accounted for before partitioning the genomic variance across the environmental variables. In this model, the spatial configuration and environmental variables respectively explained 30.1% and 8.6% of the genomic variation. However, once the adjusted r^2^ was applied to account for the number of variables, only 4.6% of genomic variation was explained by the multi-variate environment (Figure S9B; Table S4). Thus, most of the genomic variation within the eastern lineage is due to the arrangement of populations across the landscape and the serial founder phylogeographic pattern and that there is limited impact of local adaptation due to climate. To understand how this variation was distributed across the range, we predicted values for RDA axis 1 which comprised 44.5% of the constrained variation (or 2.05% of the total variation; Figure 4). Here we see that northern populations have stronger loadings as the axis is composed of high variability in monthly precipitation and low annual temperature. The Louisiana subpopulations did not show strong separation from other eastern lineage populations due to environmental variation along this axis, and tended to have moderate values among RDA axes 2-9 (Figure S10).

**Figure 4.**
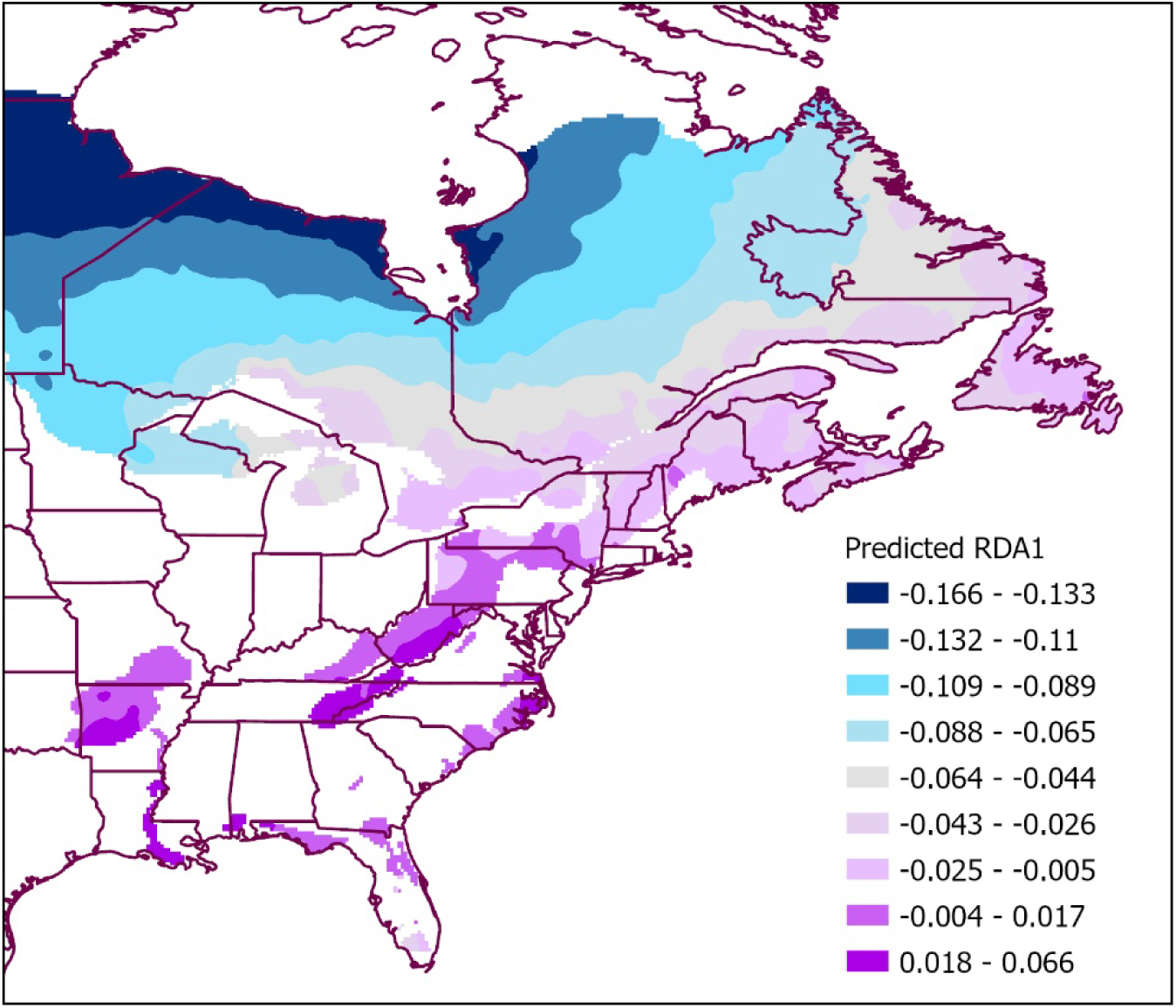
Spatial pattern of partial redundancy analysis (RDA) axis 1 across the range of the eastern lineage of *Ursus americanus* (American black bear). RDA1 explained 2.05% of the total constrained variance, whereas lineage phylogeography explained 30.1%. The axis is driven by low annual temperature and high variation in monthly precipitation in northern populations (see Figure S9B).

We used multiple approaches to consider how selection has impacted genomic variation across the eastern lineage. First, from the partial RDA, we identified sites within the tails of the first three axes. We identified 1,079 SNPs associated with the nine environmental variables; of these SNPs, 286 were annotated within 10kb of 265 genes (Table S5). Second, we conducted outlier tests among our four focal populations, including: sliding window *F_ST_* to identify highly differentiated regions, iHS to identify hard sweeps, and iHH12 to identify soft sweeps (Tables S6-S8; Figures S11-13). Across all three tests 1,653 genes were identified. We analyzed GO terms for genes with selection signatures from Louisiana (n = 597) and observed no overrepresented biological processes nor molecular function categories, reinforcing the impact of drift within these subpopulations.

To identify specific variants that may be responsible for adaptive differentiation across the lineage, we intersected regions from the outlier tests with SnpEff annotations. We identified 146 sense, 115 missense, and four nonsense variants within potentially selected regions (Table S9). Among the missense variants, 28 sites within 16 genes occurred at high frequency within one or both Louisiana subpopulations. Derived missense variants in two genes, *DHX34* and *EMCN*, occurred at high frequency in both Louisiana subpopulations. The four derived nonsense variants occurred within three genes (*CSK, DHX34,* and *PNO1*); however, none were consistently high frequency in both Louisiana subpopulations. This suggests genetic draft has had limited to no impact on the bottlenecked subpopulations of Louisiana.

### Population Genetics within Translocated Populations

Both the UARB and Missouri populations have a history of translocation (Csiki et al., 2003; Smith & Clark, 1994) and stand out as being mismatched between their geographic locations and genetic ancestry (Figures 1, S2-3). Specifically, ancestry assignments suggest that each population was replaced due to translocations from the Great Lakes in the 1960s, and within the network, these populations share more recent common ancestry. TREEMIX estimated a migration edge of 13% from the ancestral population of Tensas and Ouachita into UARB, which may represent either some Louisiana ancestry remaining or, more likely, recent admixture with the Tensas cluster as demographic recovery continues and bears expand across the LMAV (Todt et al., Accepted). The UARB population had comparable total ROH lengths to the Coastal population, although longer tracts, indicating more recent (or ongoing) inbreeding (Figure 3C).

Given the high genetic diversity of the Great Lakes population (Figure 3B) (Puckett & Davis, 2021), the potential that the translocation moved significant numbers of deleterious alleles into Louisiana is a significant concern, supported by population genomics expectations from simulation models (Kyriazis, Wayne, & Lohmueller, 2021). Thus, we quantified sharing of derived protein altering alleles among the Great Lakes, UARB, and the remaining Louisiana populations (by combining Coastal and Tensas). Very few missense and fewer LOF variants appear to have been introduced to Louisiana. The majority of the variants identified in UARB were also observed in Louisiana and Great Lakes (Figure S7A,C), demonstrating shared eastern lineage ancestry. As our counts of shared variants could display singletons in each of two populations, we further estimated allele frequencies using a 2dSFS (Figure S7B,D). While UARB was less differentiated from the Great Lakes than from the rest of Louisiana, these differences were not great for either missense or LOF variants. While some higher frequency variants were shared between the Great Lakes and UARB and between the UARB and Louisiana, alleles shared between the Great Lakes and Louisiana were at low frequencies. On the whole, shared alleles were at low frequencies in one or both populations in the pairwise comparisons, and there was limited allele sharing overall for LOF variants.

## DISCUSSION

While the cultural significance of black bears in Louisiana will always remain, our genomic data do not support the infraspecific delimitation of *U. a. luteolus* as separate from *U. a. americanus*. Using whole genome sequence data, we provide evidence at a deeper temporal resolution for patterns suggested by previous microsatellite studies: population structure and divergence between Tensas and the remainder of the eastern lineage precede the footprint of genetic drift caused by 1900s persecution (Murphy et al., 2018; Todt et al., Accepted). However, we also show that estimates of high population differentiation, both among Louisiana subpopulations and between Louisiana and the remainder of the lineage, are not the result of adaptive processes between discontinuous groups as would be expected if incipient speciation were occurring. Instead, genetic differences between groups are largely the product of severe bottlenecks and drift. Specifically, independent bottlenecks in the Tensas and Coastal populations have reduced genome-wide diversity and driven alleles to fixation. Notably, the earlier bottleneck in Tensas appears to drive patterns of population structure through strong shifts in allele frequencies via drift and purging of deleterious variants. That said, we observed load purging in both the Tensas and Coastal subpopulations, although Tensas had lower genomic load presumably from the older bottleneck allowing more time for selection to act on deleterious alleles.

We found that local adaptation has had a smaller impact than phylogeography on the distribution of genomic variation across this lineage. The strongest drivers of adaptive variation include mean annual temperature and monthly precipitation variation, where northern populations have substantial derived variation. This finding of stronger, albeit still weak, adaptive signatures in the northern range contrasts the narrative for the uniqueness of Louisiana black bear. Our analyses fully contextualize spatial-temporal patterns relevant for the conservation management of bears. Below, we outline the proposed phylogeographic path, then highlight three key landscape-level events that explain the elevated levels of genetic differentiation measured between Louisiana subpopulations from the broader eastern lineage and each other.

### Phylogeography of the Eastern Lineage of American Black Bears

The eastern lineage of black bears began separating from the western lineage approximately 138kya, with genomic data suggesting a northern route over contemporary Canada (Puckett, 2025). The patterns of population separation within the eastern lineage, namely the early divergence of the Great Lakes from southern populations, supports this interpretation. We propose that bears continued along a northern route, only moving southward when they reached the north-south mountain ranges and coastline of the continent.

The final stages of the eastern lineage expansion followed the Gulf of Mexico coastline moving westward from contemporary Florida to Texas, USA and into the Sierra Madre Oriental in Mexico (Pedersen et al., 2021). We suggest that, during this westward expansion, bears also followed the Mississippi river northwards to utilize bottomland forest resources. This hypothesis is supported by both our genomic data as well as the archeological record. Bear remains are present at archeological sites from the Late Woodland period (400 – 1100 AD) onward, suggesting that bears extended northward through the Mississippi Alluvial Plain and into the Ozark Highlands (Peles & Kassabaum, 2020). (While there is archeological evidence for bear usage in Illinois, the timing is post-colonial (Martin, 2020) and thus does not provide convincing evidence for a southwards range expansion along the Mississippi River.) Our data indicate that the modern Tensas and Coastal subpopulations accelerated divergence from each other ∼5.5kya (Figure 2A). This represents the first of several external factors impacting population connectivity and size across Louisiana. We posit that shifts in the Mississippi River between 4.5 – 7.8kya created a substantial barrier to dispersal within the LMAV (Figure 2C) and ultimately resulted in divergence into two subpopulations. Specifically, the stage 3 and 4 shifts in the valley resulted in a barrier from the eastern bluff side near Natchez southwards to the gulf coast (Gouw & Autin, 2008; Saucier, 1994). Notably, this occurred prior to the formation of the three lobes of the delta (Törnqvist et al., 1996), thus we hypothesize the ancestral range of the Coastal population was further east than the present location.

The second temporal event we observed was a severe bottleneck in the modern Tensas population around 1,000 years ago (Figure 2B). This timing coincides with substantial human-induced landscape changes in the Lower Mississippi Valley during the transition from the Late Woodland to the Early Mississippian (400 – 1100 AD) period and the progression of the Coles Creek culture (Dye, 2015; Kidder, 1998). During this time, the human footprint in the region expanded as Coles Creek human populations grew and became more hierarchically structured, interregional exchange increased, and settlements became larger and more nucleated. Concomitant land-use practices such as forest thinning, controlled burns, intensive crop cultivation, and the conversion of floodplains into agricultural fields became increasingly widespread (Abrams & Nowacki, 2008; Delcourt & Delcourt, 2004; Dye, 2015). These modifications altered forest composition, reduced mast availability, and fragmented denning habitats, which likely forced black bears into smaller, lower-quality upland areas. Archaeological evidence also shows that bears were increasingly incorporated into ritual and symbolic contexts, suggesting selective hunting pressure, while agricultural fields and other anthropogenic attractants near settlements likely prompted both opportunistic hunting and lethal control to mitigate human-bear conflict (Kassabaum & Nelson, 2016; LaDu & Funkhouser, 2019). These anthropogenic pressures were compounded by climatic stresses during the Medieval Climate Anomaly (900 – 1250 AD), which brought higher temperatures and episodic droughts to the southeastern United States (Cook, Woodhouse, Eakin, Meko, & Stahle, 2004; Stahle & Cleaveland, 1992). Collectively, these cultural and environmental factors provide a plausible explanation for the rapid decline in the Tensas black bear population around 1000 AD.

The contemporary range of the Coastal population is relatively recent, as the delta lobes that make up southern Louisiana began forming after 566 AD (Törnqvist et al., 1996). Thus, bears would have colonized southern Louisiana after sufficient plant growth was established to support their nutritional needs. We observed an independent bottleneck in the Coastal population that occurred around 1698 AD (Figure 2B). Given that this bottleneck occurred during the settlement of North America by European colonists and the start of resource extraction, we hypothesize that the combination of bear hunting and land-use change was largely responsible for this population crash. During this period, French colonists began harvesting and exporting bear skins, which included 3-5k per decade between 1700-1760 from across North America (Obbard et al., 1999), although these numbers reflect high harvest from modern Canada.

### Integrating Genomics into Conservation Policy

There is no genomic evidence supporting the subspecies level designations among the three contemporarily named groups investigated here: *americanus, floridanus,* and *luteolus*. We fully acknowledge that, without the historical context of our temporal analyses, the *F_ST_*summary statistics (Table S2) would recognize the Tensas and Coastal subpopulations as subspecies-level differentiation, even between themselves. However, we show that this is a function of genome-wide diversity loss and fixation, rather than adaptive divergence and incipient speciation.

Additional support for the subspecific delimitation of *U. a. luteolus,* such as ecological adaptation or phenotypic differentiation, is either absent or flawed. Proponents of clear criteria for subspecific delimitation emphasize that genetic differentiation alone is not sufficient: groups cannot be clinal, differences must be non-neutral, and morphological evidence (i.e., diagnosably distinct heritable phenotypes) should be present (Patten, 2015; B. L. Taylor et al., 2017). Thus, the population structure alone is insufficient for defining previously named infraspecific taxa. Furthermore, there is limited support for diagnosable phenotypes to distinguish bears among these groups. We presume that diagnosable phenotypes are encoded genomically and represent meaningful biological differentiation (Chambers & Hillis, 2019; Patten, 2015; B. L. Taylor et al., 2017), albeit without the expectation that divergence has progressed to reproductive isolation. The diagnosable phenotypes that have been put forth to distinguish *U.* (*a.*) *luteolus*, include a golden coat (Griffith, 1821); a more convex forehead and more conical nose (Griffith, 1821) with larger overall skull morphology (M. L. Kennedy, P. K. Kennedy, M. A. Bogan, & J. L. Waits, 2002; Michael L. Kennedy, Phyllis K. Kennedy, Michael A. Bogan, & Juliann L. Waits, 2002); and being “gentil in disposition” (Griffith, 1821). Each of these characteristics has limited utility as a diagnosable phenotype. First, the reports of golden bears ranging from Texas to Virginia, USA in the 1700-1800s AD is no more, as eastern lineage bear populations are greater than 95% black in coat color (Puckett et al., 2023; Rounds, 1987). We know of no historical specimens of golden bears from which to assess morphology nor genomics. It is likely that the color morph was rare on the landscape and extirpated following scientific description. Second, a single Louisiana black bear was used in the original infraspecific description, which is not a sufficient sample size by modern geometric morphometric standards to assess shape variation of a group. Discussions of this animal’s body size and pelage length ignore the time of the year in which the animal was killed, as this impacts both traits due to fat loss and gain during hibernation and hyperphagia, and spring molting, respectively. Third, the most comprehensive analysis of skull morphology within the eastern lineage included 222 bears from diverse museum collections measured 33 cranial and 11 mandible lengths (M. L. Kennedy et al., 2002; Michael L. Kennedy et al., 2002). Multivariate analysis showed clinal variation along the same range expansion axis detailed in this paper. However, allometry (i.e., shape difference due to size) was not accounted for within the analysis. Thus, it is unclear the extent that this analysis identified skull shape variation versus a skull size cline along a latitudinal gradient. We note that, while other traits such as resource use and den chronology (Fowler, Belant, Wang, & Leopold, 2019) also vary latitudinally, these traits are plastic in American black bears.

Given our results on subspecies status, concerns that have been expressed about future impacts of population connectivity creating admixture between Tensas and UARB are unwarranted (Nowak, 2025). The genomic data does support that population replacement occurred following translocation of bears from the Great Lakes region. Our data show that this translocation bridged the extremes of environmental and genomic differences within the eastern lineage; however, the overwhelming majority of the variants identified in the UARB population are shared across the eastern lineage (Figure S8). Additionally, the proportion of genetic variation determined to be locally-adapted across various regions does not seem to have had significant fitness effects for the bears translocated from Minnesota into the LMAV; the UARB currently comprises a stable subpopulation per the most recent census and population viability assessments (Clark et al., 2025). Further, there appear to be F_1_ individuals within our data (Figure S2) as bears have been able to expand out of formally isolated patches. This natural experiment has broader implications for the ability of managers to move eastern lineage bears for demographic and/or genetic rescue with limited concerns about outbreeding depression.

Regardless of subspecific status, efforts to conserve the Louisiana black bear have prevented their extirpation in the region and ongoing management will benefit these subpopulations and their ecological communities. While ESA protections have ensured demographic recovery, genomic recovery has not yet been achieved. There was a dramatic loss of genetic diversity within formally protected Louisiana subpopulations, which we show occurred due to both deeper time landscape change (Figure 2) and 19^th^ century conversions to agriculture (Figure S5). Despite evidence of purging, the low diversity, elevated levels of inbreeding, and indications of persisting isolation between some LMAV subpopulations suggests that ongoing management will be necessary for their long-term genetic health. Some of this may be achieved through protecting and restoring the bottomland hardwood forests that Louisiana bears rely upon, which would also provide critical resources for swamp rabbits, forest birds, waterfowl, and fishes.

Indications of multiple anthropogenically-driven bottlenecks in our data highlight the role of humans as a “hyper-keystone species” (Worm & Paine, 2016). Anthropogenic activities that drive bears to the point of near-eradication are not limited to the resource-extraction methods of colonial settlers or deliberate policy aimed at predator persecution, but a consequence of more general human practices: due to wide-reaching direct and indirect impacts on ecosystems, coexistence with large carnivores is a perennial challenge across human history and cultures. At present, continuing to carefully manage Louisiana bear populations will help mitigate contemporary human-bear conflict and ensure long-term preservation of valuable genetic diversity within the species.

## ACKNOWLEDGEMENTS

We thank Maria Davidson, John Hanks, and Johnny Berry (LDWF), and Dana Morin (Mississippi State University) for providing new samples from Louisiana and west Mississippi specifically for this project. Previously collected and/or sequenced samples were provided by: J. Beringer (MDC); D.R. Etter (MI DNR), K. Noyce and D. Garshelis (MN DNR); R. Cross (ME IFW); J. Hawley (CT DEEP); C.P. Carpenter (WV DNR); D. Kocha, S. Thompson, and J. Sajecki (VA DWR); C. Olfenbuttel and M. Carraway (NC WRC), and M. Orlando and M. Connelly (FL FWCC). We thank Randy Cox for discussions of fluvial change in the lower Mississippi Delta; David Dye for discussions of native peoples in the LMAV; Jay Puckett for assistance with GIS; and the University of Memphis High Performance Computer administrators for extra storage space. This work was financially supported by the National Fish and Wildlife Foundation (grant 74258) and a University of Memphis Faculty Research Grant to EEP.

## DATA ACCESSIBILITY

Sequence data has been deposited onto the NCBI SRA under project number PRJNA867575. Code is available on https://github.com/hclendenin/EasternDemography.

