## Supplemental Figures for "Population specific bottlenecks inflated differentiation measures of Louisiana black bear and negate subspecific status"

**SUPPLEMENTAL TABLE LEGENDS**

**Table S1-** List of samples used in this study. Metadata includes sample ID, US state and (sub)population where sampled, sex of the animal unless unknown (U), depth of WGS sequencing, sample name in NCBI SRA project number PRJNA867575, and diversity metrics. Diversity metrics included individual observed homozygosity (H_O_) across variable sites or genome-wide (calculated across the sum of the scaffold lengths), count of the number of Runs of Homozygosity (ROH) tracts, sum total length of ROH tracts across the genome in kilobases, and the fraction of the genome composed of ROHs (F_ROH_).

**Table S2-** Pairwise population estimates of *F_ST_* among geographic populations of eastern lineage *Ursus americanus*. *F_ST_* values are highlighted as minimal differentiation (white; 0-0.050), low (light lilac; 0.051-0.150), medium (medium purple; 0.151-0.250), and high (dark purple, >0.251).

**Table S3**- All variants (n = 43,194) within protein coding genes annotated by SnpEff within the eastern lineage of *Ursus americanus*. Position with the DNAZoo reference genome are given as scaffold and position, the SNPeff annotation is included, along with the gene name.

**Table S4-** Percentage of variance explained by each environmental variable in both the full environmental and partial redundancy analyses using ANOVA. All nine environmental variables were significant in each analysis.

**Table S5-** Variants (n = 1,079) within outlier distributions of each of the first three partial RDA axes. For each variant the environmental variable the site is most associated with is listed along with the closest gene within 10kb of the position (NA denotes there were no annotated genes within 10kb).

**Table S6-** Windows identified from 50kb sliding *F_ST_* outlier tests among four focal populations (Great Lakes, Southeast, Tensas, and Coastal) of *Ursus americanus*. Windows (scaffold, start position, and stop position) within the top 0.1% of the distribution were denoted as significant and adjacent windows were merged resulting in 220 windows across six pairwise tests. Regions were intersected with gene positions and listed (NA denotes no genes intersected with the window). If a region was also associated with a significant region identified within the redundance analysis (RDA, Table S5), then the environmental variable of the association was listed. Table is associated with Figure S10.

**Table S7-** Variants identified within single population iHS tests of the Great Lakes (GL), Southeast (SE), Tensas, and Coastal populations of *Ursus americanus* to detect hard sweeps. Selected sites comprised the 0.05% and 99.95% of the distribution of values from each test. Sites were intersected with gene positions and overlapping genes were listed (NA denotes no genes intersected with the site). Table is associated with Figure S11.

**Table S8-** Variants identified within single population iHH12 tests of the Great Lakes (GL), Southeast (SE), Tensas, and Coastal populations of *Ursus americanus* to detect soft sweeps. Selected sites comprised the top 0.1% of the distribution of values from each test. Sites were intersected with gene positions and overlapping genes were listed (NA denotes no genes intersected with the site). Table is associated with Figure S12.

**Table S9-** Intersection of genomic regions identified as outliers in any of *F_ST_*, iHS, or iHH12 tests and SNPeff annotated variants within those regions. Frequencies of the derived allele were calculated for the four focal populations: Great Lakes, Appalachian Mountains, Coastal, and Tensas.

**SUPPLEMENTAL FIGURES**

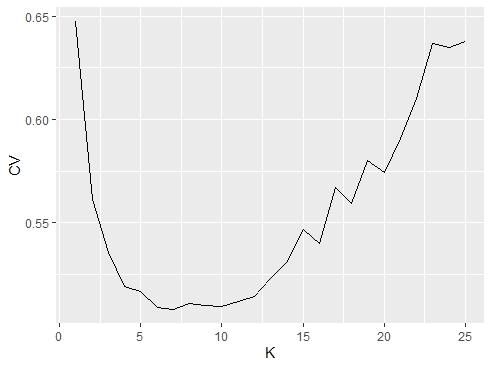

**Figure S1-** Cross validation error (CV) plots from ADMIXTURE for the eastern lineage of *Ursus americanus* (American black bears). Clustering was run from 1 to 25 clusters (K) for 20 repetitions of the program to estimate cross validation error.

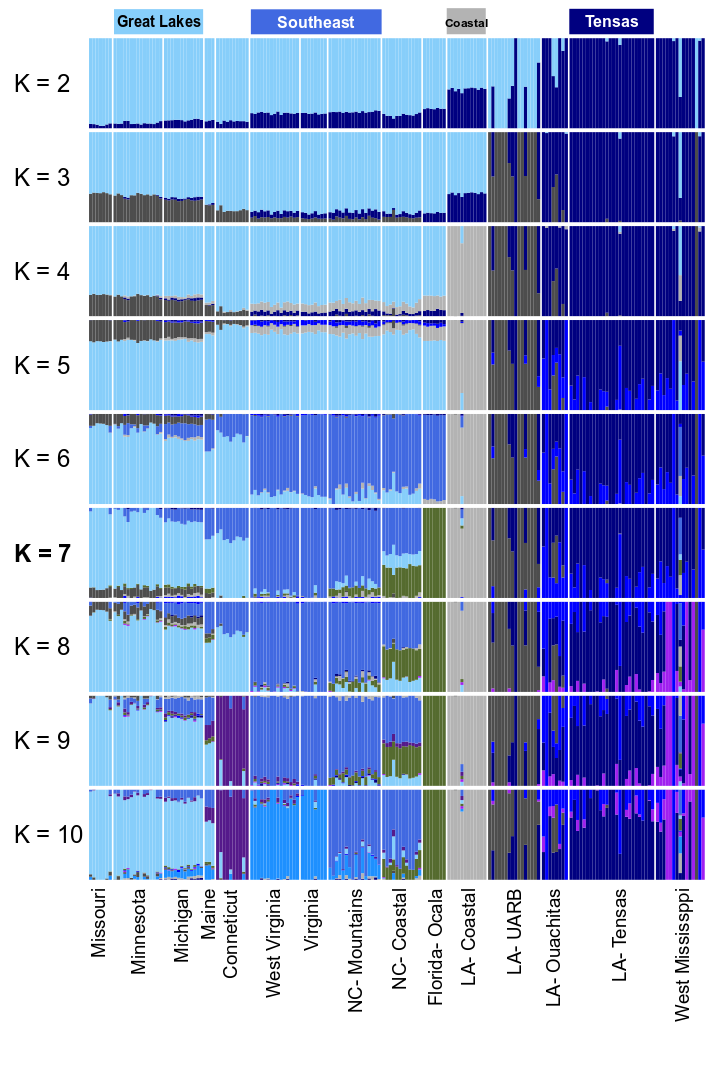

**Figure S2-** Population structure of *Ursus americanus* (American black bear) across 183 samples from eastern North America as estimated by ADMIXTURE. Ancestry proportions from seven clusters (K = 7) represents the best supported model (see Figure S1). The four focal populations (Tensas, Coastal, Southeast, and Great Lakes) discussed heavily in the main text are highlighted above the admixture plots, while USA state names and/or geographies are listed below.

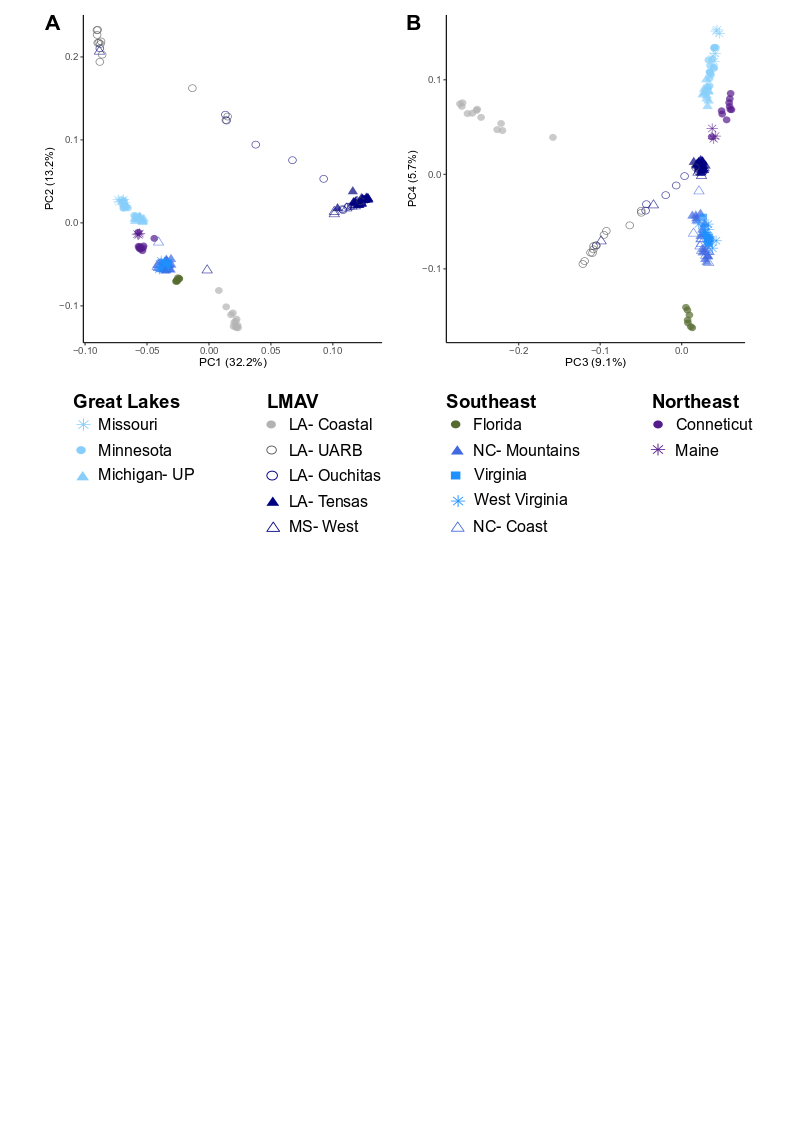

**Figure S3-** Principal components analysis of *Ursus americanus* (American black bears) eastern lineage samples. (A) Axis 1 (32.2% of the variation) separates the Tensas subpopulation in Louisiana (navy triangles) from most other eastern lineage samples. Axis 2 (13.2% of the variation) represents a latitudinal gradient with the exception of the Upper Atchafalaya River Basin (UARB) subpopulation in Louisiana (open grey circles) which is the replacement population following translocation of bears from the Great Lakes region in the 1960s. (B) Axis 3 (9.1%) separates the Coastal, Louisiana population from the remaining eastern lineage diversity. Notably, UARB samples which appear as fully replaced with the Great Lakes signature on PC axis 1 and in ADMIXTURE results, load more negatively than other populations on this axis. Axis 4 (5.7%) broadly separates the main range expansion axis similarly to axis 1, although with limited inference of Louisiana samples. Samples are denoted by color and symbology based upon their collection location.

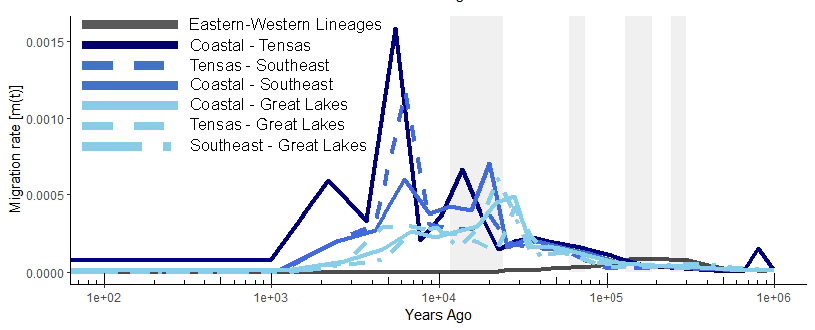

**Figure S4-** Bidirectional migration rates [m(t)] estimated for eastern lineage populations of *Ursus americanus* (American black bear) using MSMC-IM. Colors and line styles indicate population pairs, including: Great Lakes and Coastal (light blue and solid); Great Lakes and Tensas (light blue and regular dash); Great Lakes and Southeast (light blue and long-short dash); Southeast and Coastal (medium blue and solid); Southeast and Tensas (medium blue and regular dash); and Coastal and Tensas (navy and solid). As a reference point a comparison between the eastern and western lineages was included as a dark grey and solid line. Light grey background bars indicate glacial periods identified as Marine Isotope Stages (MIS) 2, 4, 6, and 8 (left to right).

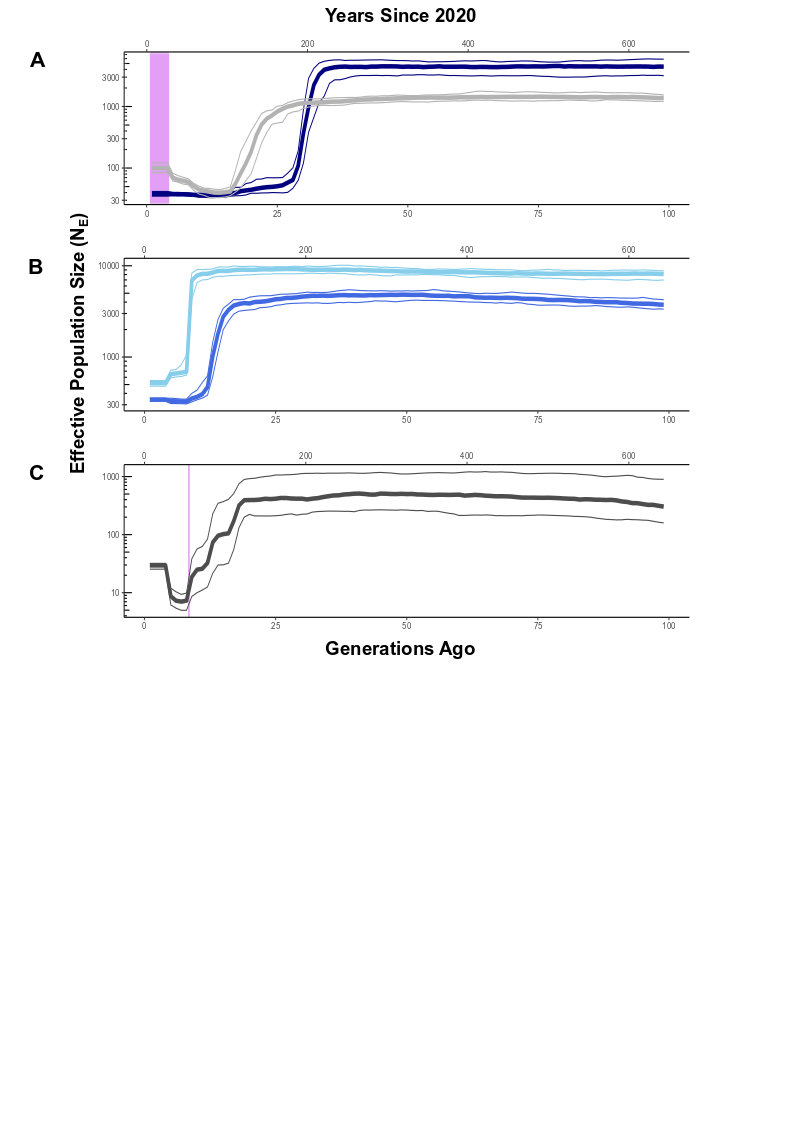

**Figure S5-** Recent change in effective population size (N_E_) across the eastern lineage of *Ursus americanus* (American black bear) estimated using GONE. Thick lines represent the median of 50 repetitions, while thin lines are the 2.5% and 97.5% confidence intervals. We use 2020 as the reference year from which to estimate timings of size change in years and a generation time of 6.5 years per generation to covert the top axis of each plot. (A) The Tensas (navy) and Coastal (light grey) populations in Louisiana show sharp N_E_ declines occurred 20-30 generations ago. The violet box highlights the duration in which these subpopulations were listed under the US Endangered Species Act. (B) Populations from the Great Lakes (light blue) and Southeast (medium blue) indicate modern populations crashes between 8-15 generations ago. (C) Change in N_E_ within the contemporary Upper Atchafalaya River Basin (UARB) subpopulation in Louisiana that shows population replacement by Great Lakes genotypes since a translocation of bears between 1964-1967 (violet vertical line).

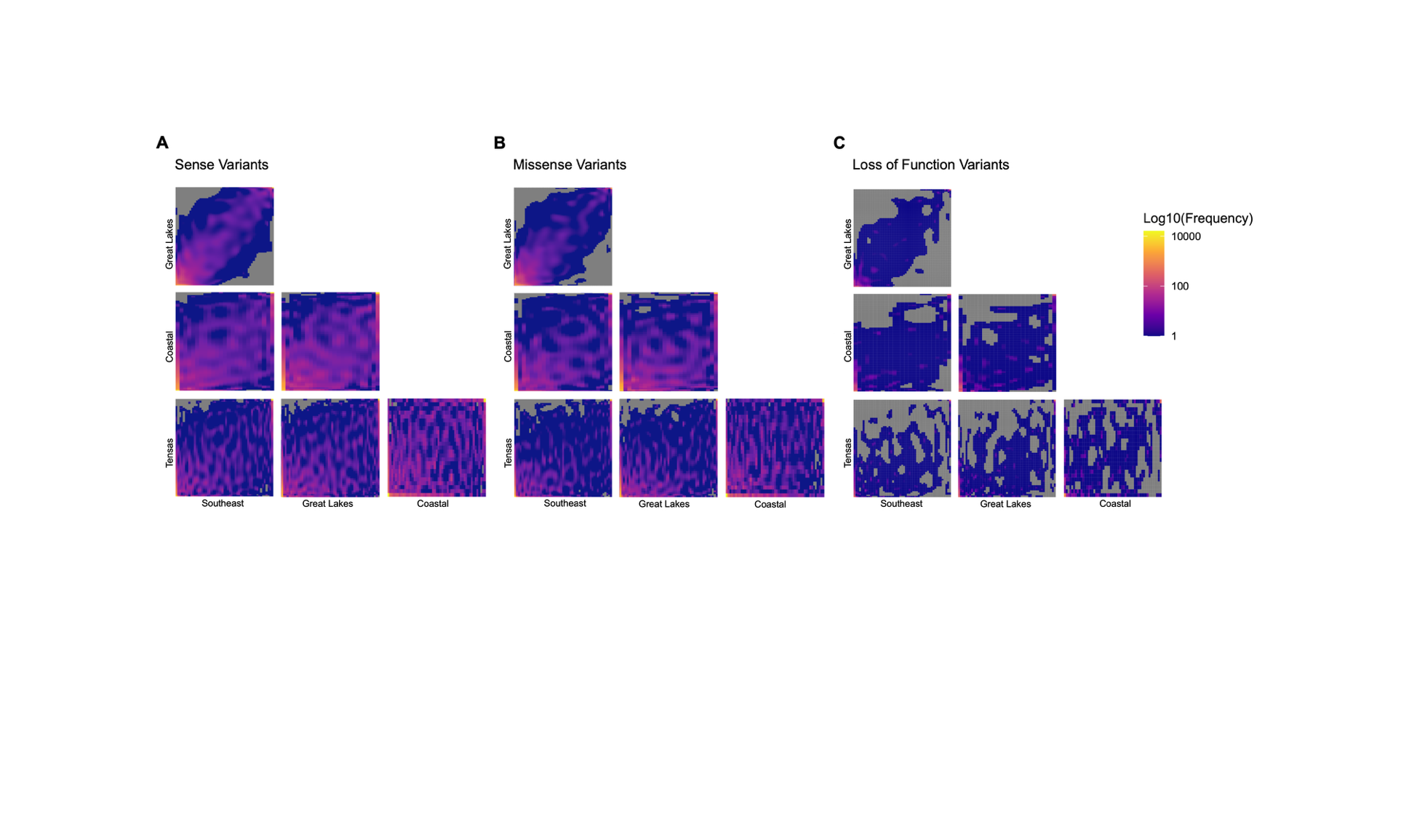

**Figure S6-** Allele sharing among four focal populations (Great Lakes, Southeast, Coastal, and Tensas) of the eastern lineage of *Ursus americanus* (American black bear). Pairwise two-dimensional site frequency spectra (2dSFS) were visualized where the count of unique derived alleles is represented by the cell color across the full matrix of chromosome counts per population. The scale was log 10 transformed to highlight differentiation on the low end of total variation. Cells with no shared alleles are grey. Plots were made among sense (A), missense (B) and loss of function (C) variants as annotated by SnpEff.

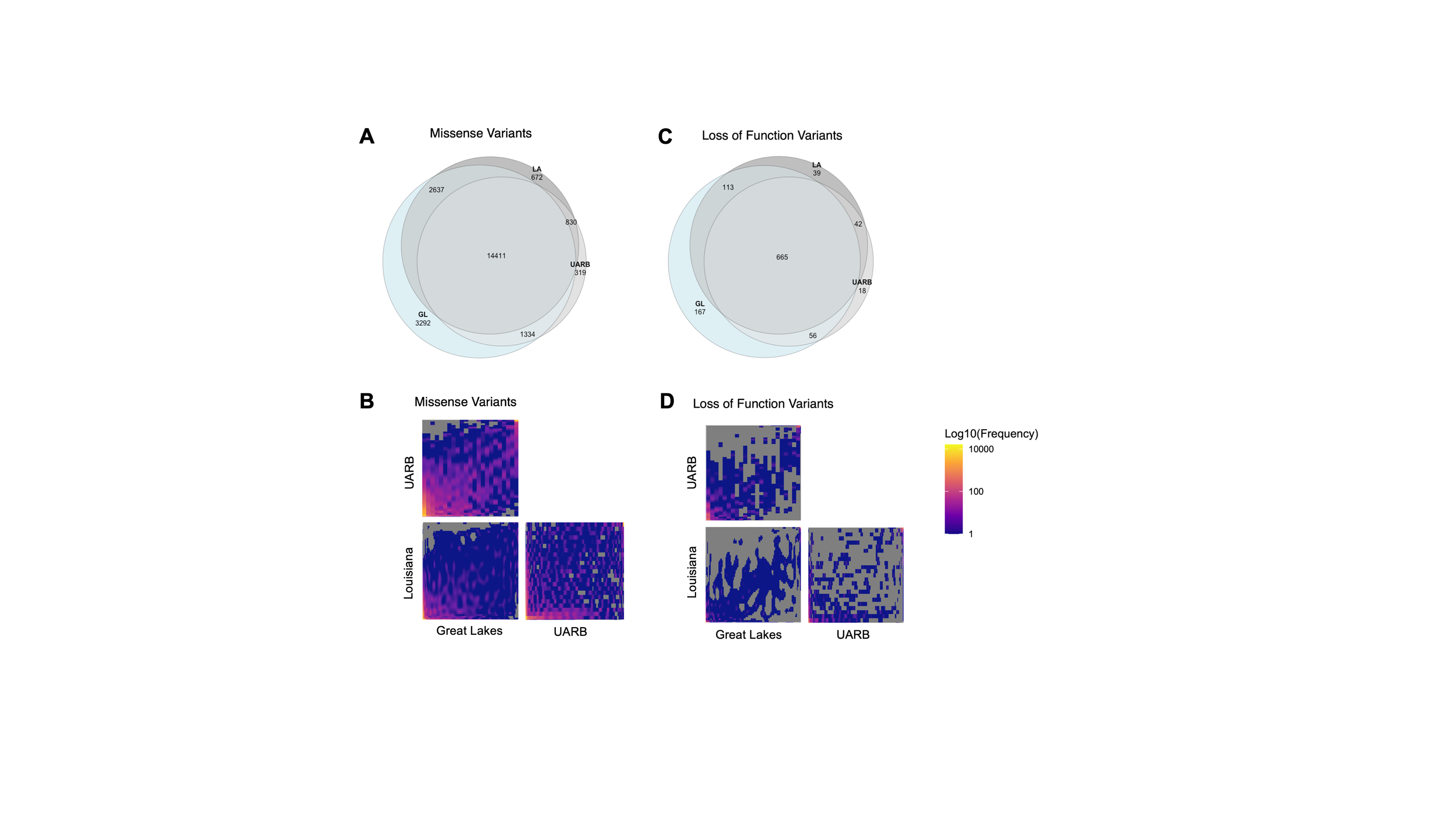

**Figure S7-** Allele sharing of missense (A-B) and loss of function (C-D) variants among three populations (Great Lakes, a combined Coastal and Tensas, and the contemporary Upper Atchafalaya River Basin [UARB]) of the eastern lineage of *Ursus americanus* (American black bear). (A, C) Euler plots were generated from counts of absolute shared and private sites. However, (B, D) two-dimensional site frequency spectra (2dSFS) were calculated to understand the impact of population-level allele frequencies. Counts of derived alleles are represented by the cell color across the full matrix of chromosome counts per population. The scale was log 10 transformed to highlight differentiation on the low end of total variation. Cells with no shared alleles are grey. Individuals with less than 90% UARB ancestry (see Figures 1C, S2) were removed prior to calculations.

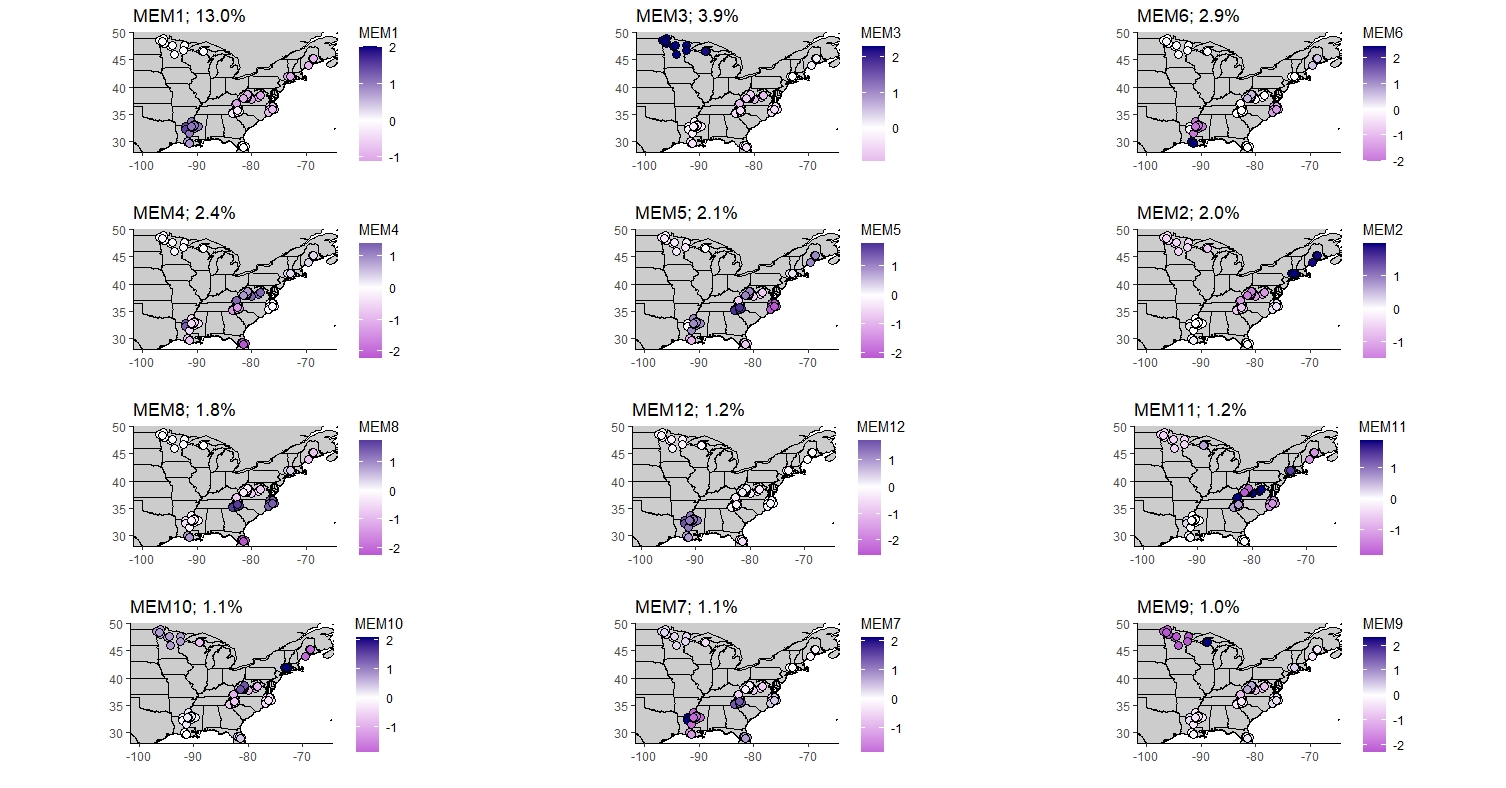

**Figure S8-** The 12 significant distance-based Moran’s Eigenvector Maps (dbMEMs) used within the redundancy analysis (RDA) to account for the spatial configuration of the samples across the landscape. Eigenvectors are organized from the most significant to the least (moving left to right, top to bottom). Note that MEM3 was removed prior to the full and partial RDA analyses due to high correlation with multiple BioClim variables. The proportion of genomic variation that each MEM explains is listed above the map.

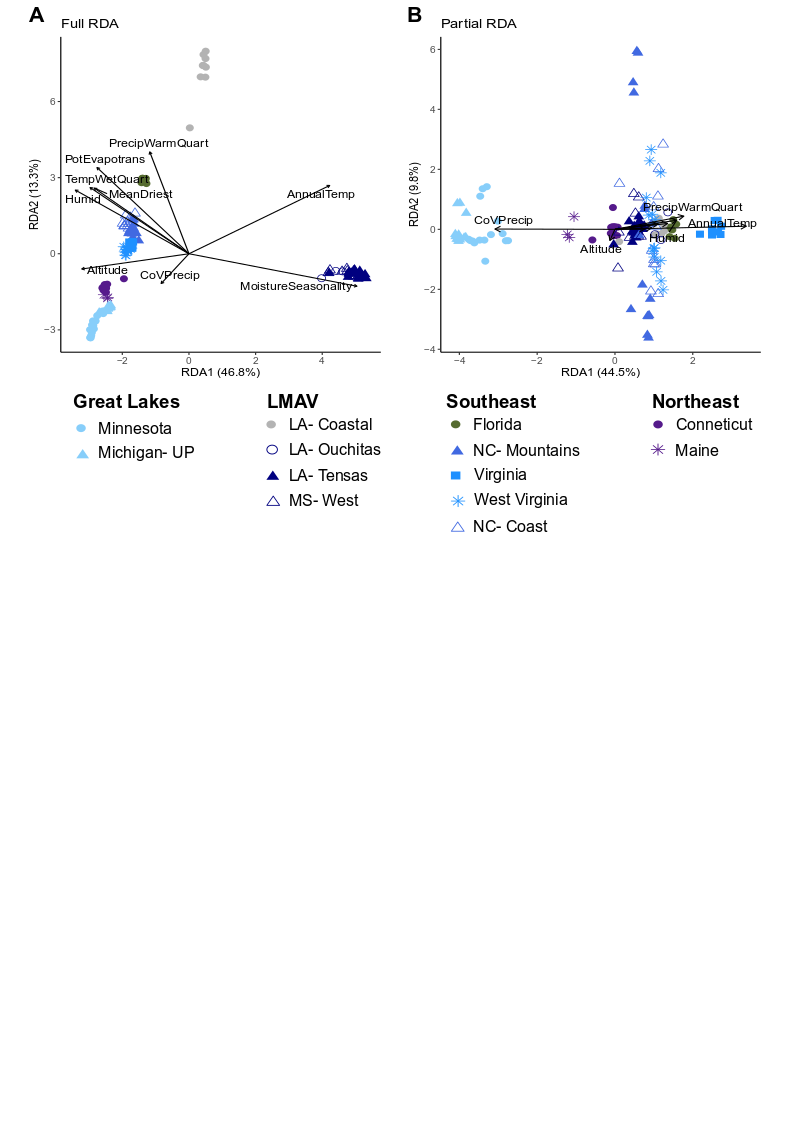

**Figure S9-** Redundancy analysis (RDA) biplots from the (A) full and (B) partial models which, respectively, did not and did account for the variance attributed to the spatial configuration of populations among the eastern lineage of *Ursus americanus* (American black bear). Arrow lengths of tested environmental variables were multiplied by 10 so they would be visible within plots, indicating weaker contributions to the variance than in the display.

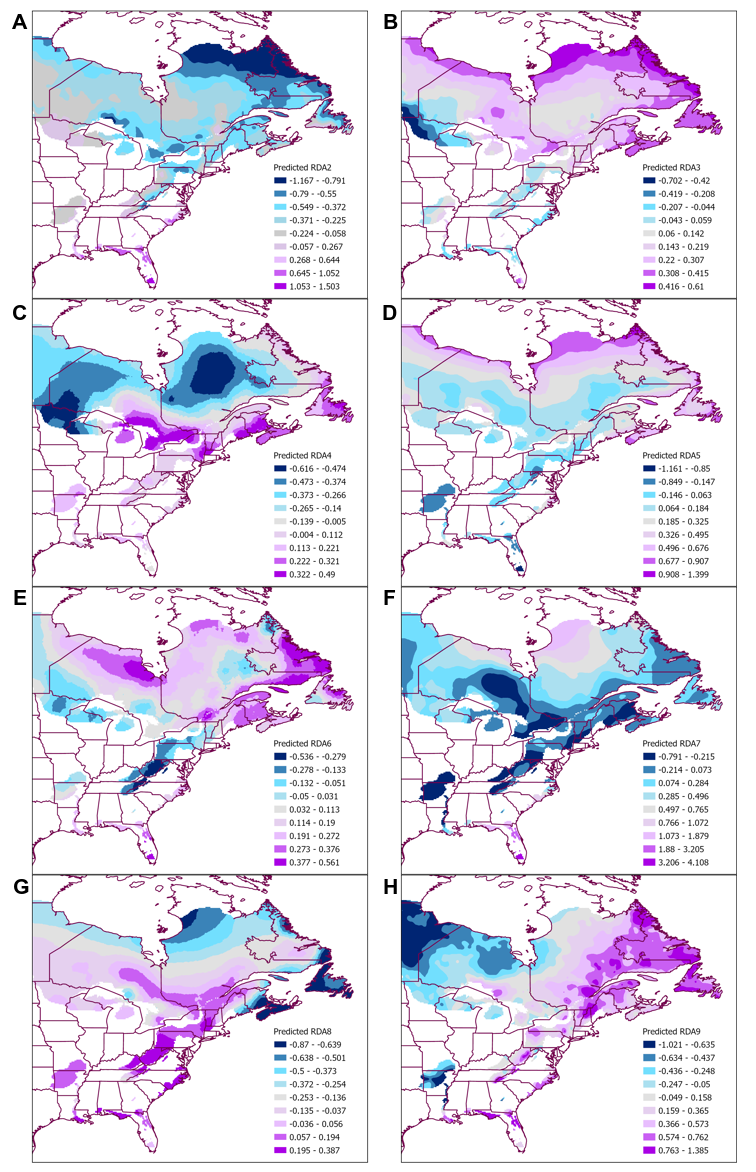

**Figure S10-** Predicted and spatially smoothed redundance analysis (RDA) axes 2-9 (panels A through H, respectively) across the eastern lineage of *Ursus americanus* (American black bear). Values in the legend represent the range of the predicted axis as divided via nine Jenks natural breaks.

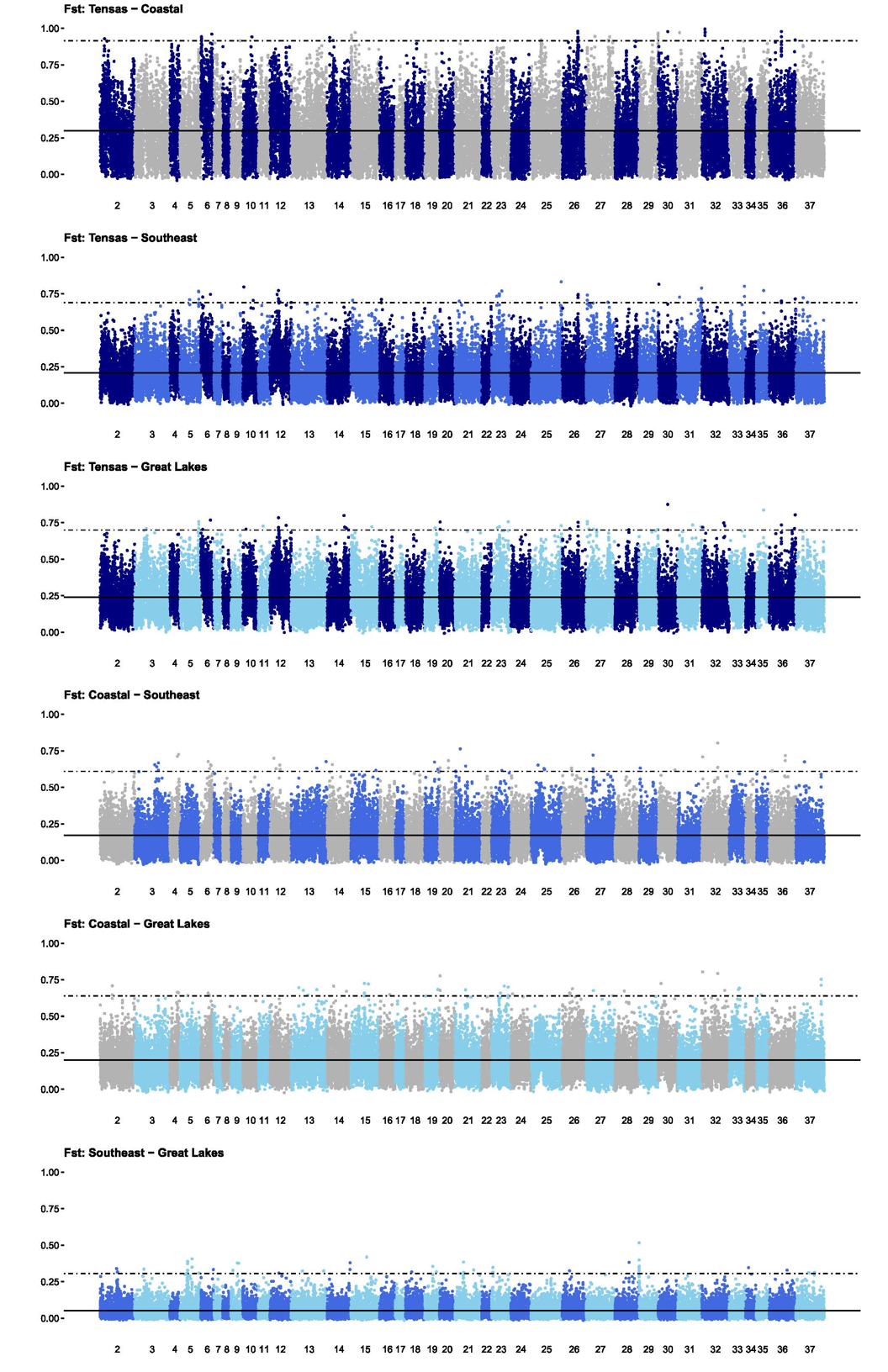

**Figure S11-** Differentiation (*F_ST_*) estimated pairwise among four focal populations (Great Lakes- light blue, Southeast- medium blue, Coastal- grey, and Tensas- navy) of the eastern lineage of *Ursus americanus* (American black bear). Within each panel the solid line represents the genome-wide average *F_ST_* between the two populations, while the dashed line shows the 99.9^th^ percentile above which sites were considered possibly to be selected. Figure is associated with Table S6.

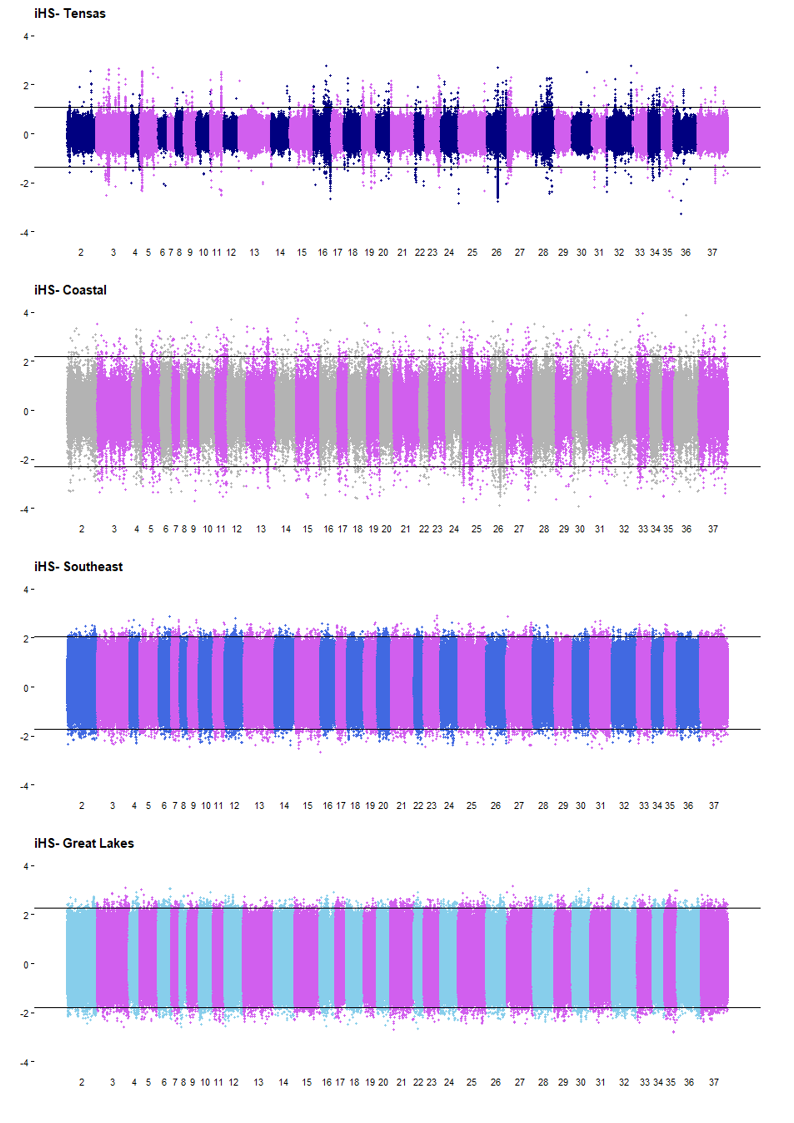

**Figure S12-** Genome-wide integrative haplotype scores (iHS) to test for hard sweeps pairwise among four focal populations (top to bottom: Tensas- navy, Coastal- grey, Southeast- medium blue, and Great Lakes- light blue) of the eastern lineage of *Ursus americanus* (American black bear). iHS was calculated in sliding windows (20kb window and slide) across scaffolds 2-37 using selscan. The horizontal lines represent iHS scores of the 0.5^th^ and 99.5^th^ percentile, for which windows respectively below or above were considered possible regions of selection. Figure is associated with Table S7.

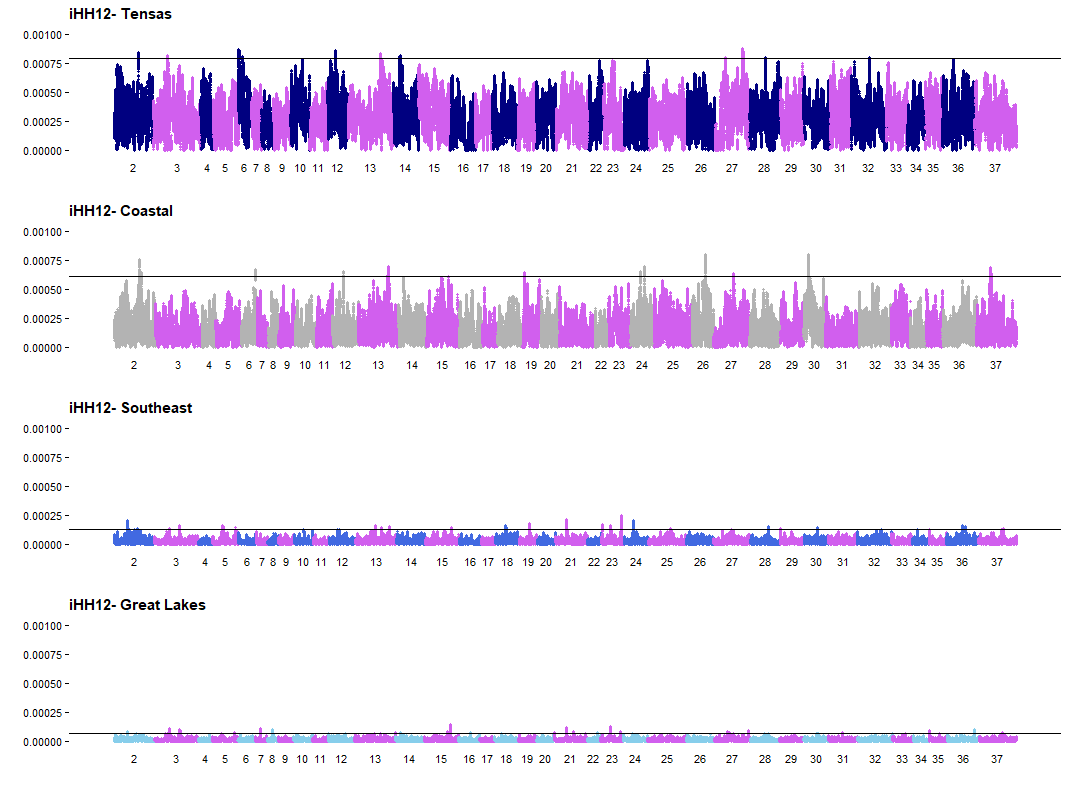

**Figure S13-** Genome-wide integrative haplotype homozygosity pooled (iHH12) to test for soft sweeps within each of four focal populations (top to bottom: Tensas- navy, Coastal- grey, Southeast- medium blue, and Great Lakes- light blue) of the eastern lineage of *Ursus americanus* (American black bear). iHH12 was calculated in sliding windows (20kb window and slide) across scaffolds 2-37 and normalized by allele frequency using selscan. The horizontal line represents the per population 99.9^th^ percentile, above which windows were considered possible regions of selection. Figure is associated with Table S8.
